# Modeling control of invasive fire ants by gene drive

**DOI:** 10.1101/2025.03.04.641467

**Authors:** Yiran Liu, Samuel E. Champer, Benjamin C. Haller, Jackson Champer

## Abstract

*Solenopsis invicta*, commonly known as the fire ant, is characterized by aggressive behavior and exceptional invasive capabilities, rendering conventional control methods largely ineffective. Here, we explore the implementation of homing suppression gene drive in fire ants by developing a spatially explicit model that incorporates both monogyne and polygyne colony structures, enabling comprehensive evaluation of genetic control strategies. Ants may present unique challenges for gene drive due to their colony structure and haplodiploidy. Our results reveal that after an extended period of time, gene drive effectively eliminates polygyne colonies, but monogyne populations often persist at low level. Though standard suppression drives in haplodiploids have reduced power, new dominant-sterile resistance or two-target strategies, as well as drives that affect the colony structure, can restore high suppressive capability. Interspecific competition can also exert a positive effect on gene drive-mediated population suppression dynamics. In particular, a gene drive release during the invasion phase significantly enhances population suppression, enabling native ants to successfully recolonize their original habitats. We also identify several conserved female fertility genes in fire ants, together with gRNA targets that may support efficient, low-resistance suppression drive designs. Overall, we conclude that while gene drive in fire ants may take place over extended time scales, its long-term results, even with imperfect efficiency, are quite promising.

## Introduction

The invasive fire ant, *Solenopsis invicta*, is a haplodiploid species notorious for its remarkable invasive capacity and resistance to control efforts. Accidentally introduced nearly a century ago from the Pantanal region in South America^1^, fire ants have spread to numerous countries worldwide, including the United States^2–4^, China^5^, Australia^6,7^, and others^8^. Recently, they were reported in Italy in 2023^9^, marking the first recorded presence of this formidable species on the European continent. Damage caused by fire ants is extensive, encompassing ecological disruptions such as declines in native biodiversity^10,11^, agricultural loss through crop destruction^12,13^, and public health concerns due to their venomous stings and aggressive behavior^14–16^. Various strategies have been explored for controlling fire ant populations. Bait traps have demonstrated some effectiveness in eliminating fire ant colonies, but their use poses risks to wildlife and domestic animals^17^. Biological control involving pathogens and parasites have also been investigated^18^. For example, an endoparasite nematode showed promising efficacy in the lab, but its performance was less effective in the field^18^. Such challenges, including pesticide resistance, ecological side effects, and limited efficacy, highlight the ongoing difficulty in identifying a reliable and sustainable approach for managing fire ant populations.

Gene drive technology is a unique approach for managing agricultural pests or disease vectors. By biasing inheritance in its favor, a gene drive can increase in frequency within a population^19–21^. By targeting fertility or viability genes, suppression gene drive systems can ultimately lead to the eventual elimination of the population. To date, suppression gene drives have been successfully implemented in the laboratory in various organisms, including fruit flies^22^, spotted wing drosophila^23^, mosquitoes^24^, and medflies^25^.

Despite the growing interest in utilizing gene drive to manage invasive species^26^, the application of this approach to fire ant suppression remains unconsidered. Fire ants are a haplodiploid species, meaning that males are haploid and possess only one set of chromosomes from their mother, while females, which develop from fertilized eggs, are diploid. Haplodiploid species exhibit enhanced resilience to inbreeding depression due to their sex determination system^27^, which both diminishes the suppressive effects of this phenomenon and increases the probability of resistance evolution in females^28^. Previous studies have demonstrated that homing gene drive still has the potential to be an effective tool for controlling haplodiploid pests, though efficiency is lower than in diploids^29,30^. However, the unique lifecycle of fire ants presents additional challenges for the feasibility of gene drive applications. Specifically, the complex interactions between genetic inheritance and colony structure in fire ants may influence the success of gene drive strategies.

Colonies typically consist of a queen, female workers, and fertile ants (also named alates).

Alates engage in nuptial flights, after which mated queens subsequently establish new colonies. These colonies are mostly stationary and can persist for several years. Females sterilized by a suppression drive could not produce workers and would thus not be competitive, potentially slowing suppression. Fire ant colonies can be classified into two distinct social forms. Those with a single reproductive queen are referred to as the monogyne form. Polygyne colonies have more than one queen. Monogyne and polygyne colonies substantially differ in behavioral ecology. For example, the queen in monogyne colony, on average, has a longer life span than the smaller queens of polygyne colonies^31^. Polygyne queens prefer to choose males in or near their own nest, and the colony reproduces by budding, while monogyne queens find males through nuptial flights, allowing for wider dispersal^32^. Moreover, polygyne colonies exhibit reduced aggression toward other colonies with the polygyne social form^33^. These social forms are closely associated with a “greenbeard” gene at a single genetic locus, which contains two alleles, *SB* and *Sb*, the latter of which yields a dominant polygyne phenotype^34,35^. Polygyne social forms manifest greater ant densities than monogyne colonies in many places (despite individually smaller workers), resulting in increased ecological perturbation and economic costs^36^.

Numerous modeling studies on suppression gene drive systems have revealed unexpected complexities when applied to spatially structured populations. These challenges often arise from the result of uneven suppression across the geographic distribution^37^, which influences the spread of the drive and persistence of wild-type^38^. Environmental heterogeneity adds another layer of complexity, as variations in niche carrying capacity and local resource availability can influence the survival and reproductive success of individuals^39,40^. Together with other spatial dynamics^28,41^, these can significantly influence the propagation and effectiveness of gene drives and potentially lead to their failure. On the other hand, interactions with other species may also impact the outcome of gene drive strategies. In particular, the presence of competing or predator species could contribute to target species suppression^40^.

To determine the potential of gene drive for fire ant suppression, we model a homing suppression drive targeting a female fertility gene within a two-dimensional spatial framework including fire ant lifecycle characteristics. To enhance suppression outcomes, we also consider several improved drive variants. These include dominant-sterile resistance^42^, a two-target drive design^43^, and hypothetical drive systems that affect fire ant social form.

Recognizing that fire ants are an invasive species, our model also incorporates native ant populations to evaluate the potential role of native species in mitigating fire ant spread in conjunction with a gene drive. We found that while the complete suppression of fire ant populations required an extended time period, polygyne colonies were eliminated more rapidly. Greater suppression could be achieved with some of our improved drive strategies.

Furthermore, the presence of native ant species also facilitated the suppression of fire ants, and releasing drive individuals during the invasion process could further accelerate target population elimination. Bioinformatic analysis shows that several promising female fertility target genes are highly conserved in fire ants and amenable to a gRNA multiplexing strategy. Overall, our findings provide a greater understanding of suppression gene drives in invasive fire ants and highlight the potential of utilizing native biodiversity in biocontrol strategies.

## Methods

### Suppression drive strategy

Fire ants, as a haplodiploid species, have different chromosomal patterns than more common diploids. The egg-laying queens are diploid, while the males only have one chromosome, developing from unfertilized eggs. Accordingly, we model a homing suppression drive targeting a haplosufficient but essential female fertility gene, which is the only type of powerful, self-sustaining suppression drive that has shown to be viable for haplodiploids^30^. The gene drive mechanism is implemented for each individual queen (the type of drive has no activity in haploid males). In drive/wild-type heterozygotes, the wild-type allele is cleaved by CRISPR/Cas9, which is specifically guided by one or more guide RNAs (gRNAs). The cleaved chromosome then undergoes homology-directed repair, which results the drive allele being copied to the wild-type site (“drive conversion”). Female drive homozygotes will be sterile. With the growing fraction of sterile females as drive frequency increases, the population size will be decreased or even eliminated eventually if the fraction reaches a high enough level (based on the power of the drive).

If conversion does not occur, the wild-type allele may instead mutate into a resistance allele, a process referred to as germline resistance allele formation. Such resistance alleles cannot be recognized and cut by Cas9. Resistance may also arise when offspring inherit a wild-type allele while the mother carries a drive allele. In such cases, any wild-type allele in the offspring may be converted into a resistance allele due to the activity of maternal deposited Cas9 and gRNA during early embryonic development (we refer to this as the embryo resistance allele formation rate).

Resistance alleles are usually nonfunctional (referring to disruption of the female fertility target gene). This kind of resistance still could cause female sterility but will slow down the drive process and reduce its power, especially if the drive has fitness costs. In another less frequent but more drastic outcome, the mutation will cause suppression drive to fail almost immediately if the resistance allele preserves the function of the target gene. In such scenarios, females retain reproductive capacity due to the functional resistance allele, which will gain a selective advantage and rapidly outcompete the drive. However, experimental evidence demonstrates that targeting a highly conserved gene together with gRNA multiplexing can mitigate the emergence of functional resistance^24,44,45^. A synergistic combination of these offers a promising approach to prevent functional resistance formation. Hence, we only consider nonfunctional resistance in this model. However, we previously found that haplodiploid species are somewhat more susceptible to the emergence of functional resistance alleles than diploids^30^, reinforcing the need for careful experimental design for gene drives in haplodiploids.

Drive female heterozygotes may experience fitness costs due to unintended Cas9 activity (leakiness) in somatic cells. This off-target activity disrupts wild-type alleles in somatic cells, resulting in reduced fertility among female drive heterozygotes. This heterozygote fitness cost may also be caused by haploinsufficiency of target genes or even arise from desired germline activity, depending on the expression pattern of the female fertility target gene or cleavage of the wild-type allele in other ovary cells by Cas9.

### Fire ant reproduction model

The forward-in-time genetic simulation framework SLiM (version 4.3) was used for our model^46^. In the fire ant model (Figure 1), all colonies are represented by a single queen, which has a genotype for herself and a genotype for her previous mate. The model assumes a default population size of 50,000 monogyne colonies and 30,941 polygyne colonies (so that each social form occupies 50% of territory at the start, see below), randomly distributed within a 1.85 km × 1.85 km territory (which usually can support 100,000 monogyne colonies at equilibrium in the model). Time steps are yearly, which represents a full reproductive cycle in fire ants.

**Figure 1.**
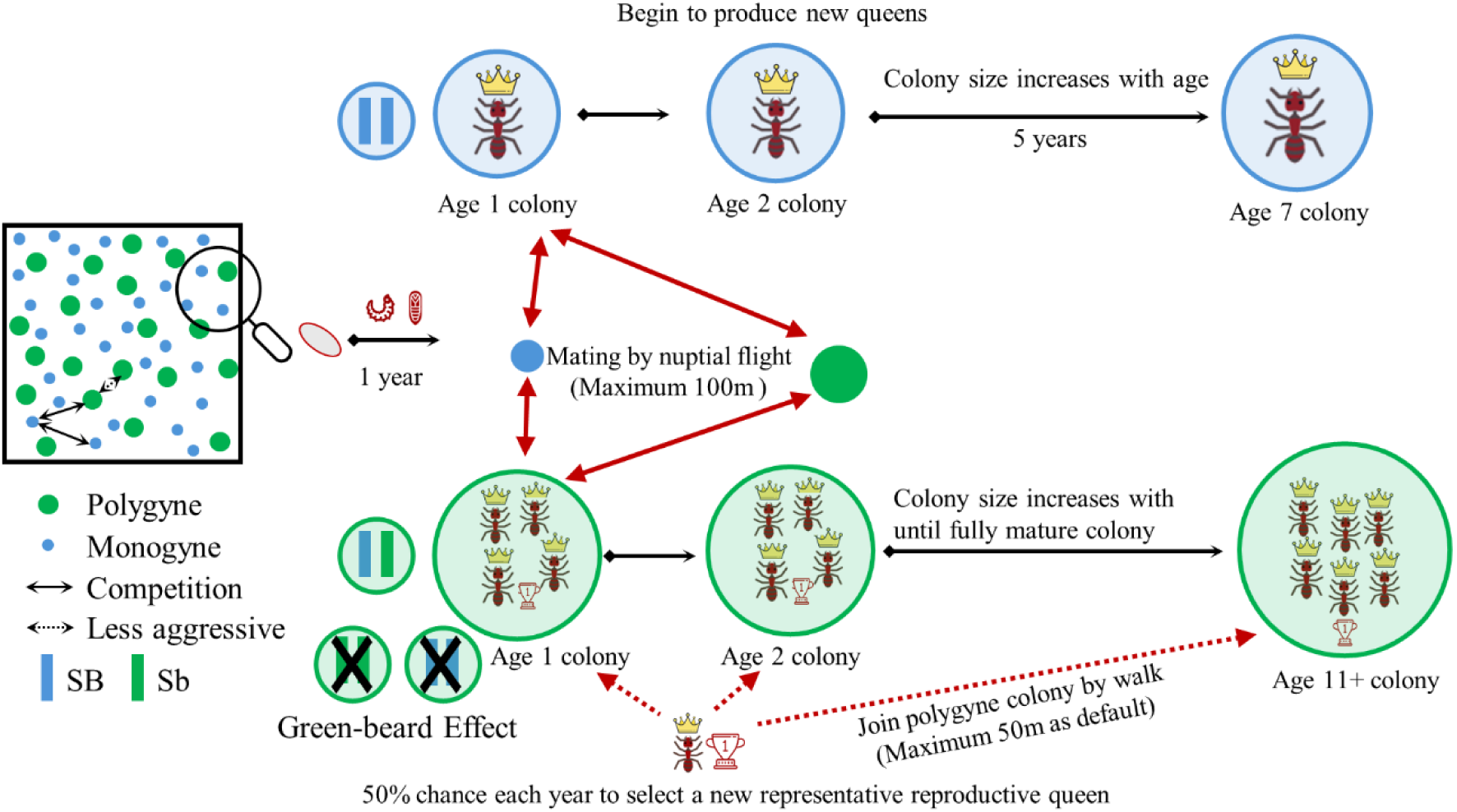
Diagram of the fire ant model. Two types of fire ant social forms are randomly distributed within the territory. Green dots represent polygyne colonies, while blue dots indicate monogyne colonies. Black arrows in the diagram illustrate competition between colonies, with dashed arrows representing weaker competition. The colonies are influenced by the greenbeard effect and Sb homozygote nonviability, meaning that polygyne colonies only give rise to SB/Sb heterozygous females. Red arrows indicate the nuptial flight and the distance for establishing a new colony, while red dashed arrows represent the shorter-ranged expansion of polygyne colonies through budding.

Only colonies older than age 1 are capable of reproduction. In the program, age 1 queens can mate (selecting a mate from a colony capable of producing offspring) but cannot produce any offspring until age 2. Note that fire ants mate before establishing a colony, but this takes place after the first year of density-dependent competition in our model to produce a nearly identical result with lower computational burden. This is possible because age 0 colonies are too small to exert any significant competition on their neighbors. We assume that age 1 new queens have up to 10 attempts to find a neighboring colony from which to take a mate, with the fitness of potential colonies determined by their genotype. Larger colonies also have a proportionately higher (based on their biomass) likelihood of being selected. Furthermore, we reduce the rate that polygyne colonies are selected by the “polygyne male rate” because polygyne colonies are known to produce fewer males^47,48^. This is set to 0.837352526 by default so that the higher biomass and slightly different lifecycle of polygyne colonies does not allow them to be selected at a higher rate.

The number of new queen progeny is generated based on a Poisson distribution and is based on the age of that colony. This is because even though polygyne colonies are larger, the workers are typically smaller and less healthy than those in monogyne colonies. Polygyne colonies may be less healthy per unit of biomass, but the new queens they produce are smaller too. We thus approximated that both colonies produce the same number of queens at the same age, though drive queens produce proportionally fewer based on their fitness cost. Specifically, we assume that 10,000 ants are needed for basic colony maintenance, and that each additional 10,800 ants in the colony produces a single new queen (the actual number of female alates produced is larger, but we reduce this number to reduce the computational burden while still allowing high variation in the survival of offspring from each colony, even at low density when survival is high).

After the offspring are produced, they can migrate based on a normal distribution to build their own colony around their mother with a standard deviation equal to *average dispersal*∗*√ π* / 2. The position is redrawn if the offspring is placed outside the boundaries of the arena. Our default monogyne average dispersal distance is 100m (0.054159 in SLiM)^49^, while the dispersal distance for polygyne colonies is set at half of this value due to lack of sufficient body mass/energy reserves to support sustained flight^50^.

Age 0 colonies have a survival of:

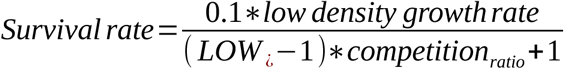

The biological differences between monogyne and polygyne ant colonies have implications for colony behavior, social dynamics, reproductive strategies and so on. Polygyne colonies are composed of multiple queens with a shorter lifespan than monogyne queens^35,51,52^. Further, newly mated polygyne queens typically join existing polygyne colonies rather than establish their own colony^53,54^. To simplify our model, we still represent this colony as single genotype, but we allow it to change its genotype more rapidly than the lifespan of a normal polygyne queen, representing the influx of new queens that increasing dominate offspring production. Thus, each year, all polygyne colonies have a chance to replace their representative reproductive queen genotype, which we set as 50% as default. Colonies unsuccessful in queen replacement experience colony mortality with a probability of 80%. If they remain viable, the genotype of the previous queen persists. Further, the greenbeard effect in polygyne colonies, associated with the acceptance of multiple polygyne queens within a nest, results in *SB/SB* genotype new alate queens from polygyne colonies being eliminated by workers^55^. The *Sb/Sb* genotype is generally lethal in females^55^, so polygyne colonies only produce *SB*/*Sb* (polygyne) queens and tend to have 50% female reproductive individual mortality (the *SB*/*SB* and *Sb*/*Sb* new queens). It is not known exactly what may happen to monogyne colonies with offspring carrying polygyne alleles. To make monogyne alleles somewhat competitive (though still at a frequency-dependent disadvantage, see below) with polygyne alleles, we assume that new monogyne queens will only mate with *SB* males, even if their mate is from a polygyne colony.

### Colony mortality

The survival rate of fire ant colonies in our model is influenced by their age and the level of competition they experience from other colonies. Monogyne fire ants queens can live up to 7-8 years^56^. Thus, we implement a 50% age-related death rate of age 6 colonies, and no colony survives past 7 years. Limited knowledge is available about the lifespan of polygyne colonies, which are characterized by multiple queens and continual influx of new queens.

Thus, we assume that they can persist for up to 70 years (chosen because the number of colonies that reach this age is negligible).

Following the observed growth rate of new colonies^57^, we increase the colony size as it ages, which affects how much competition that colony exert on others and how much competition will affect their own mortality rate. The colony size growth curve is approximately logistical^58^. Monogyne colonies follow the equation: 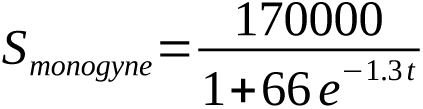

where *S* is the colony size (in number of ants) and *t* represents the age in years. Polygyne colonies usually contain ∼2 times the number of workers as monogyne colonies of similar age^36^, so here we used the same colony size equation form but changed the maximum colony size to 340,000: 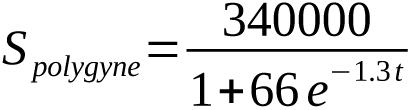

Competition experienced by each colony is calculated as:

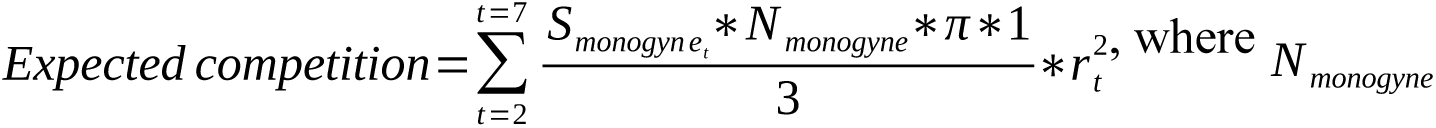 represents the total carrying capacities for monogyne-only populations, with default values of 100,000. *S_t_* means the colony size in *t* year, and *r_t_* represents the competition radius in *t* year separately. The factor of 1/3 comes from the average interaction strength between the colony that received the competition and the colonies that is the source of the competition, which linearly declines with distance. The competition radius of a monogyne colony is based on territory size.

Mature colonies have a territory of 100 m^2^, and younger colonies have a proportional territory area based on their biomass relative to the mature colony. The radius yielding this territory size is then doubled, to allow colonies with overlapping territory to compete. Polygyne colonies are assumed to have the same territory area as monogyne colonies of the same age.

Fire ant colonies will compete for resources and often directly battle^59^. Thus, we implement density-dependent mortality for all colonies. with monogyne colonies having a density-dependent survival rate of:

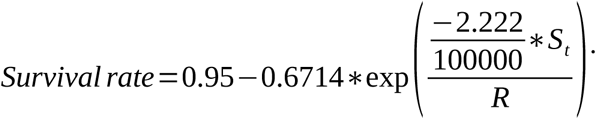

The competition ratio (R) in equation refers to *competition ratio* (*R*)=*actual competition* / *expected competition.* Actual competition experienced by each focal colony is determined by the cumulative competitive pressure exerted by neighboring colonies of both social forms (monogyne and polygyne). In spatial models this competition acting over the course of several years tends to produce colonies that are substantially more dispersed compared to a random distribution. Thus, to maintain desired growth rates and carrying capacity, the number of offspring was multiplied by 0.95, which was empirically determined to within 1%.

Polygyne colonies have the same survival rates as monogyne, except that the first factor of 0.95 is replaced by 0.9. This is because despite their increased size, multiple polygyne colony queens still have greater total maintenance burdens than a single monogyne queen. Further, polygyne queens tend to be smaller and less healthy (lower survival rates), and some queens only produce sterile alates, reducing colony efficiency^31,32,56^. Polygyne colonies exhibit reduced average worker size relative to monogyne colonies (workers average approximately 60% the size of a monogyne worker)^60^, resulting in diminished competitive capacity. To account for this phenotypic variation in our model, we apply a scaling factor of 0.6 to represent the reduced competitive ability. Thus, the higher number of ants in polygyne colonies still allows them to exert 20% more competition on their neighbors than monogyne colonies of the same age. Additionally, polygyne colonies do not attack other polygyne colonies^61,62^, resulting in reduced competition between them. Of course, they still consume resources that are thereby denied to other nearby polygyne colonies, competing indirectly. We incorporate this by implementing an intraspecific competition coefficient of 1/1.2 between polygyne colonies in the actual competition calculation (the exact resource and energy usage here has not been studied, and we thus assume that larger polygyne colonies exert the same amount of competition pressure on each other as a smaller monogyne colony of the same age). Overall, this allows polygyne colonies to slowly outcompete monogyne colonies in the course of a simulation with default parameters in absence of gene drive intervention (Figure S1). This appears to match field observations^63,64^, though the greater dispersal ability of monogyne allows it to remain ecologically viable in real-world settings where additional factors may be present^65^.

### New drive variants

We also incorporated two updated homing suppression strategies into our model. The first is the dominant sterile resistance strategy, in which the presence of a single resistance allele leads to female sterility^42^, and females lacking a wild-type allele are also still sterile. Though conceivable based on our previous study, the population dynamics of such a drive in haplodiploids or diploids has not been assessed to our knowledge. Another strategy is the distant-site/two-target suppression drive, where the homing drive targets and rescues an essential gene at its own site, with additional gRNAs inducing cleavage at a separate female fertility gene^43^. This strategy often had improved genetic load (suppressive power) in diploids compared to standard suppression gene drive, depending on the exact performance parameters, but has not been assessed in haplodiploids.

Given the shorter generation time of polygyne colonies, we modeled novel gene drives associated with this social form to investigate the impact on suppression gene drive efficacy. Three variants of our standard homing suppression drive were considered.

Polygyne drive: This drive contains a copy of the *Sb* gene, which determines the fire ant colony social form. If a monogyne female inherits one copy of the drive, it will develop into a polygyne colony and adhere to all the rules that apply to polygyne colonies (see below).

Colonies will remain viable if they are *Sb*/*SB* heterozygotes with a drive.

Greenbeard drive: This drive rescues *SB* homozygote new queens from the greenbeard effect in polygyne colonies (where they are normally removed). They will be able to create new drive monogyne colonies.

Cleave *SB/Sb* drive: In this drive, in addition to its normal functions, it can also cleave the *SB* allele in the germline of polygyne queens, converting it into a *Sb* allele. Thus, all new queens will survive in polygyne colonies where the queen mated with a wild-type male. However, if the queen mated with a polygyne male, all workers would be *Sb* homozygotes, and the queen will thus be treated as sterile/nonviable. We also modeled the reverse situation where Sb alleles are converted to SB alleles in new queens. For both drives, we assume that this process happens with ideal efficiency.

### Native ants

To evaluate the impact of native ants on the effectiveness of homing suppression drive in controlling fire ants, we introduced another ant species. In this model, native ants are represented solely by a single species with monogyne colonies. To simplify the model, we assume that both species share the same life cycle and reproduction rules. However, the competition strength experienced by each colony comes not only from its own species but also from the competing species. Hence, the expected competition in each species is calculated as:

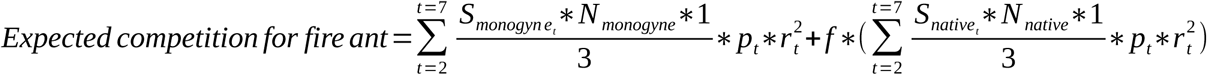

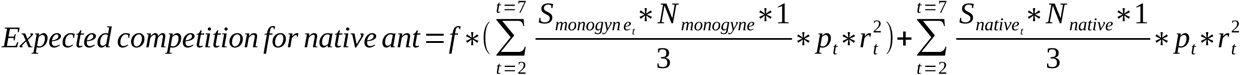

, where *f* represents an interspecies competition factor between fire ant species and native ant species with a value between 0 and 1. Thus, each species has a reduced interspecific competition compared to intraspecific competition. In the absence of gene drive, both species would thus exist at equilibrium. The default capacity is set at 50,000 for monogyne fire ant colonies, 30,941 for polygyne fire ant colonies, and 50,000 for native ants, with all other parameters remaining consistent with the fire ant-specific model. Thus, native ants are assumed reach only half the total territory size of fire ants in the region, with the underlying assumption that they are only able to persist in certain microhabitats or ecological niches that do not fully overlap with fire ants.

### Model scenarios

For most simulations, gene drive males equivalent to 15% of the age 1 colony population were released randomly over the entire arena, of which half had the polygyne *Sb* allele. This release was repeated for the first six years of the simulation.

We also modeled the invasion process of fire ants. For these simulations, fire ants were initially limited to the left 40% of the modeling arena, with the left half of this region occupied by polygyne form fire ants at equilibrium density, while monogyne fire ant colonies were similarly placed in the right half of this region. The remaining 60% of the arena was filled by native ants. Given the highly invasive nature of fire ant species, the competition factor between the two species was set to 0.5, but in this scenario, this only reduced the competition exerted by native ants on fire ants. In contrast, native ants experienced full competition pressure from fire ants. This represents a scenario in which native ants would eventually be driven to extinction by fire ants in the modeled region. The average dispersal distance in this model was somewhat increased to 138.75m to allow for more rapid fire ant invasion.

Unless otherwise specified, simulations are ended 100 years after the drive release if the population or the drive were not eliminated earlier. Default values for parameters (which are used unless otherwise specified) are shown in the Table 1. The default drive parameters were calibrated based on a successfully constructed drive in *Anopheles gambiae*^66^ representing an effective drive, but still with several imperfections. Population dynamics were quantified using an ant biomass metric^67^, with units defined as the mass of a monogyne colony worker:

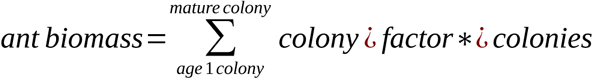

**Table 1.**
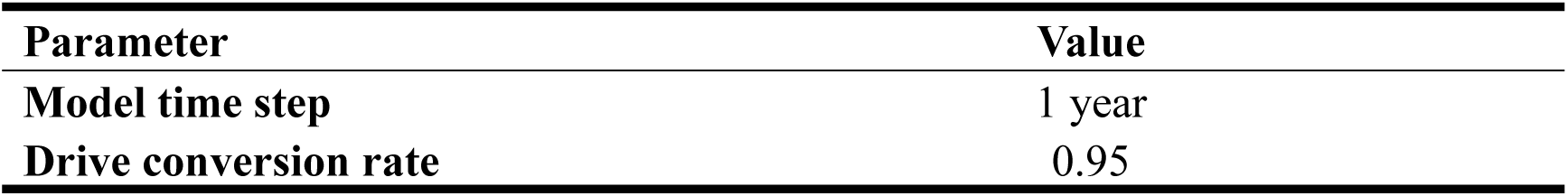
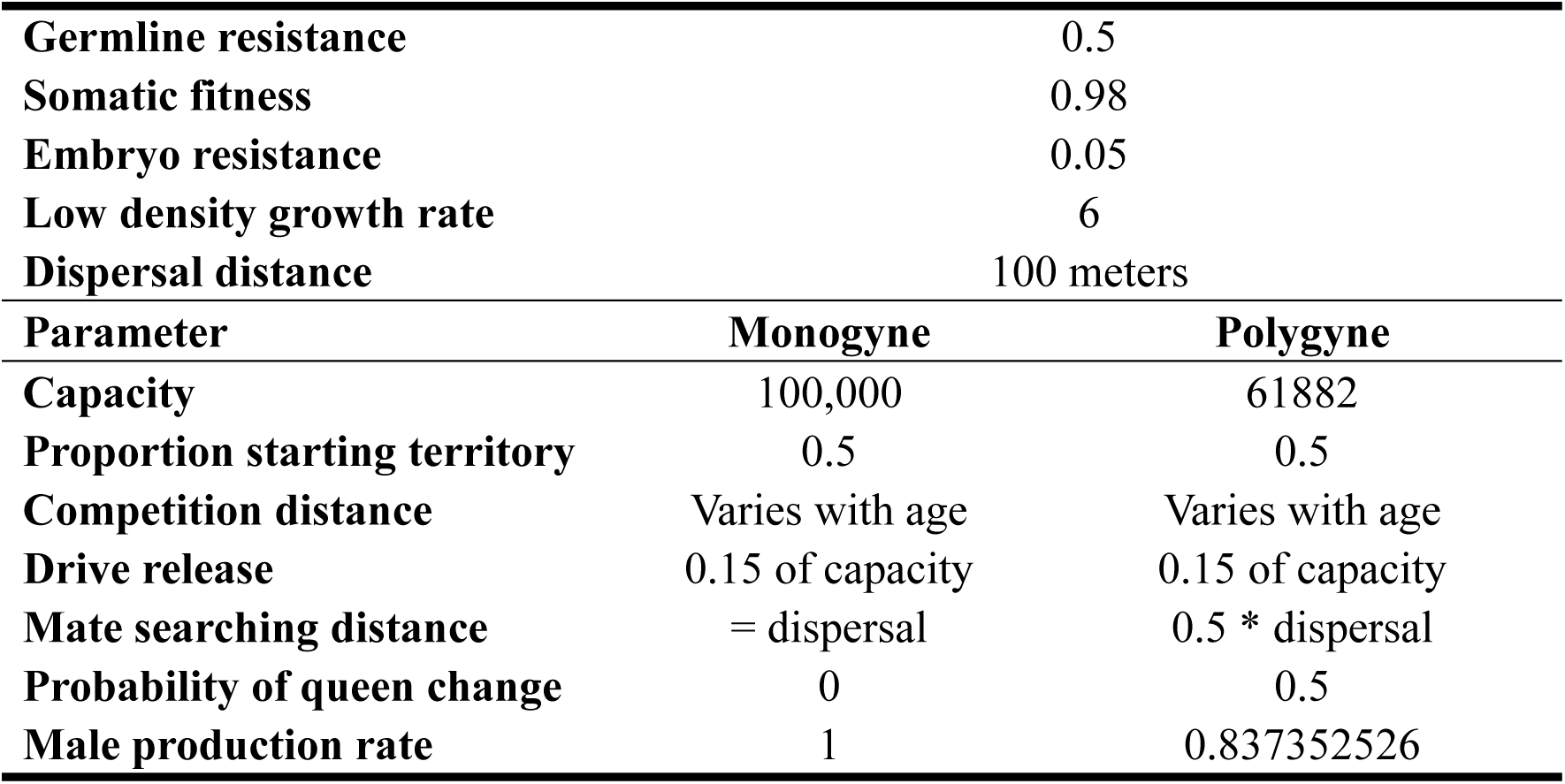
Default model parameters.

| Parameter | Value |
| --- | --- |
| Model time step | 1 year |
| Drive conversion rate | 0.95 |

|  |  |  |
| --- | --- | --- |
| **Germline resistance** | 0.5 |  |
| **Somatic fitness** | 0.98 |  |
| **Embryo resistance** | 0.05 |  |
| **Low density growth rate** | 6 |  |
| **Dispersal distance** | 100 meters |  |
| **Parameter** | **Monogyne** | **Polygyne** |
| **Capacity** | 100,000 | 61882 |
| **Proportion starting territory** | 0.5 | 0.5 |
| **Competition distance** | Varies with age | Varies with age |
| **Drive release** | 0.15 of capacity | 0.15 of capacity |
| **Mate searching distance** | = dispersal | 0.5 * dispersal |
| **Probability of queen change** | 0 | 0.5 |
| **Male production rate** | 1 | 0.837352526 |

Given the phenotypic distinctions between social forms, where polygyne colonies exhibit larger colony size but produce smaller workers with reduced fitness compared to monogyne colonies^60^, we use a mass factor of 1 for monogyne colonies and 0.6 for polygyne colonies (based on the relative worker size). We often use biomass output as a proxy for overall effectiveness of drive strategies.

### Analysis of potential target genes for fire ant homing suppression gene drive

Protein and nucleotide sequences were obtained from the NCBI database (https://www.ncbi.nlm.nih.gov/). Multiple sequence alignments were performed using the MUSCLE algorism^68^, and the resulting alignments were visualized using the platform Jalview version 2^69^. gRNA target sites were identified using the online tool CHOPCHOP (http://chopchop.cbu.uib.no).

### Data generation

All simulations were conducted on the High-Performance Computing Platform of the Center for Life Science at Peking University. Python was used for analyzing data and preparing the figures. All models and data can be accessed on GitHub (https://github.com/jchamper/Fire-Ant-Suppression-Model).

## Results

### Suppression gene drive in fire ants

To evaluate the effectiveness of a suppression gene drive in fire ants, we evaluated a homing suppression gene drive in our spatial model. Fire ant default parameters are based on estimates and field studies (see methods). We first examined the drive dynamics in a monogyne population, a polygyne population, and a mixed population (Figure 2). The drive frequency increased in all scenarios, indicating successful propagation of the drive (Figure 2A). Following drive introduction, both ant biomass (number of monogyne workers or equivalent mass, Figure 2B) and colony abundance (Figure 2C) declined. While polygyne colonies exhibited complete elimination, monogyne colonies persisted by the end of the simulation (Figure 2B-C). This was likely mainly due to the difference in generation time (Figure S2). In a hypothetical panmictic fire ant population at equilibrium, the generation time of monogyne colonies in our model is 5.4 years. For polygyne colonies, with 50% representative queen replacement per year (due to higher individual queen mortality), the generation time is expected to be slightly higher than 2 years (drive allele frequencies matched exactly at 2.3 years, see Figure S2). Though spatial and density-related factors could influence these results, polygyne ants should in general still have a substantially shorter generation time, likely accounting for most of the difference in suppression time.

**Figure 2.**
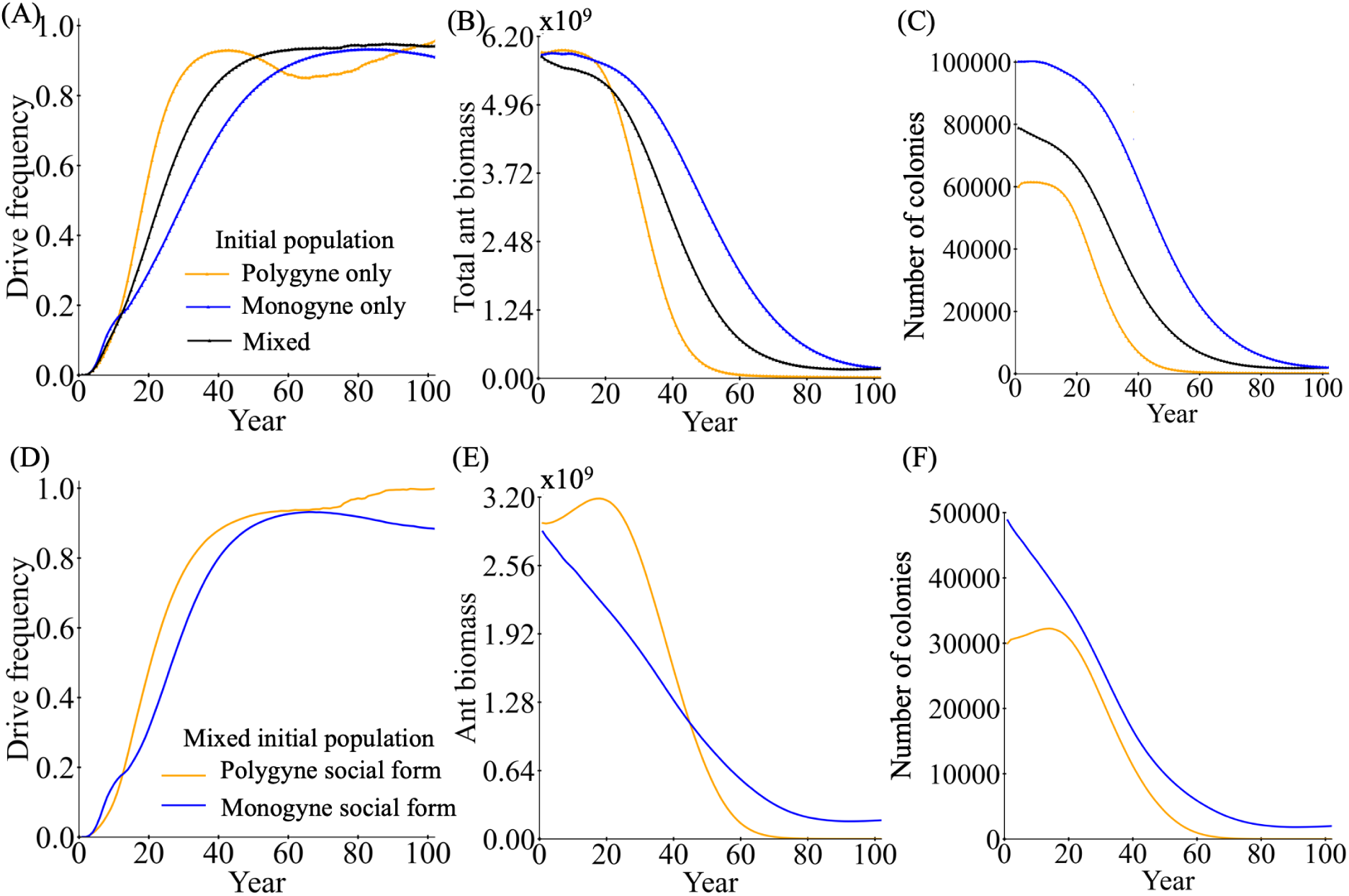
Homing suppression gene drive in fire ant populations. Drive males were introduced into the spatial fire ant population. We examined fire ant populations that were composed only of monogyne colonies, only polygyne colonies, or a mix of both, with each occupying 50% of the initial territory. We display the (**A**) drive frequency, (**B**) total fire ant biomass, and (**C**) number of colonies in each model. For the mixed population, we also separate track the (**D**) Drive frequency, (**E**) ant biomass, and (**F**) number of colonies for each social form. Each line shows the average of 200 simulations.

We also examined individual monogyne and polygyne trajectories in a mixed population. Though polygyne can outcompete monogyne without gene drive, the polygyne population was still eliminated more quickly than the monogyne population, even though monogyne experienced a more rapid initial decline (Figure 2D-F). Persistence of monogyne colonies at the very end of the simulation (despite the slower population reduction) for the mixed population was likely due to a chasing effect because our drive with default parameters theoretically had enough genetic load (0.87) to eliminate the population with a low-density growth rate of 6 (which requires a genetic load of 0.83 in a deterministic model).

Due to the long generation time of fire ants, increased importance should be placed on larger initial release sizes compared to other gene drive applications. Varying the release size, we found that modest benefits can be obtained with larger releases (Figure S3), though these decrease rapidly much past our default release rate of 15% males for six years.

### Evaluating the effect of model parameters on population suppression

We next evaluated how drive performance affects the suppression outcome. In this analysis, we varied the drive conversion and female fitness parameters from 0.8 to 1.0, representing a theoretically efficient drive capable of suppressing generic haplodiploid panmictic populations over most of the range^70^. Under a perfect drive scenario, nearly the entire fire ant population could be eradicated (Figure 3A-B). Even when fitness and drive conversion rates were reduced, the drive maintained a significant impact, decreasing the total ant biomass by more than 50% (Figure 3A). However, even with a perfect drive, achieving a 90% reduction in ant biomass required approximately 30 years (Figure 3C). Notably, the impact of reduced somatic fitness was minimal when the drive conversion rate was sufficiently high, underscoring the importance of drive efficiency in determining suppression outcomes.

**Figure 3.**
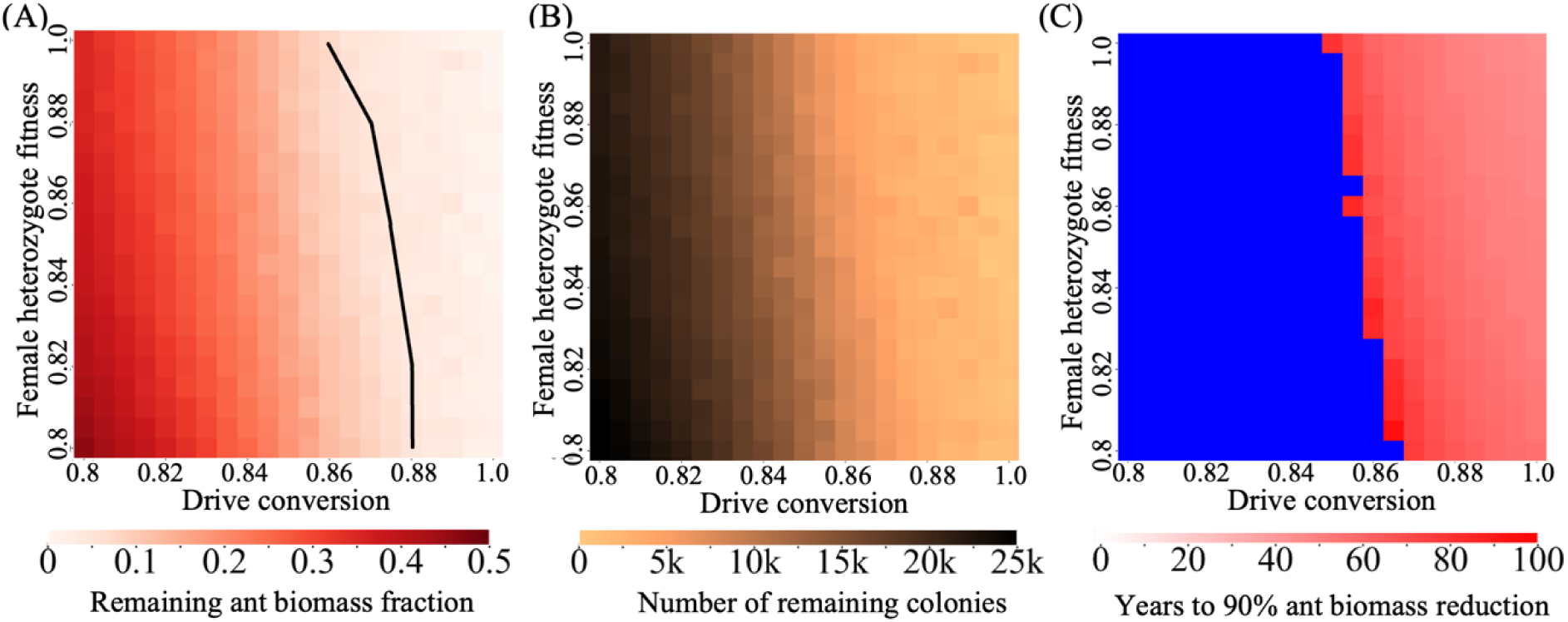
Impact of varying drive efficiency and somatic fitness on fire ant control. The drive, with varying drive conversion and female fitness, was released into a mixed spatial fire ant population. Heatmaps show the (**A**) fraction of remaining fire ant biomass compared to the initial biomass after 100 years, (**B**) the number of colonies remaining in the arena 100 years after the release of drive males (the initial number is 80,941), and (**C**) time required to achieve a 90% reduction in fire ant biomass. Blue indicates that the population fails to reach a 90% reduction within 100 years after drive release. The black line in panel A (and area to the right of the line) shows where the drive has sufficient theoretical genetic load value at equilibrium to eliminate a panmictic deterministic population. Each point shows the average result from 20 simulations (including only simulations that reached 90% biomass reduction).

The habitat occupied by fire ants and the extent of their dispersal may significantly influence the effectiveness of control efforts. To investigate these factors, we varied the growth rate of fire ants in low-density regions and their average dispersal distance (Figure 4A-C). Our findings indicate that increased dispersal distance can contribute to a subtle reduction in the fire ant population. However, its impact remains minimal when the low-density growth rate is sufficiently high. This may be because higher dispersal tends to inhibit chasing, which is a phenomenon where a suppression drive fails to eliminate the target population in a spatial model, despite theoretically having enough suppression power. Specifically, it involves wild-type individuals escaping to empty space and gaining reproductive advantages due to reduced competition. Even though the gene drive remains present, population elimination can be indefinitely delayed. Such chasing has occurred in many spatial models of suppression gene drive^28,70–72^. Chasing may play a role in late stage population suppression in our fire ant model (see below). Of course, chasing only tends to be significant when the genetic load (suppressive power) of the drive is theoretically capable of eliminating the target population.

**Figure 4.**
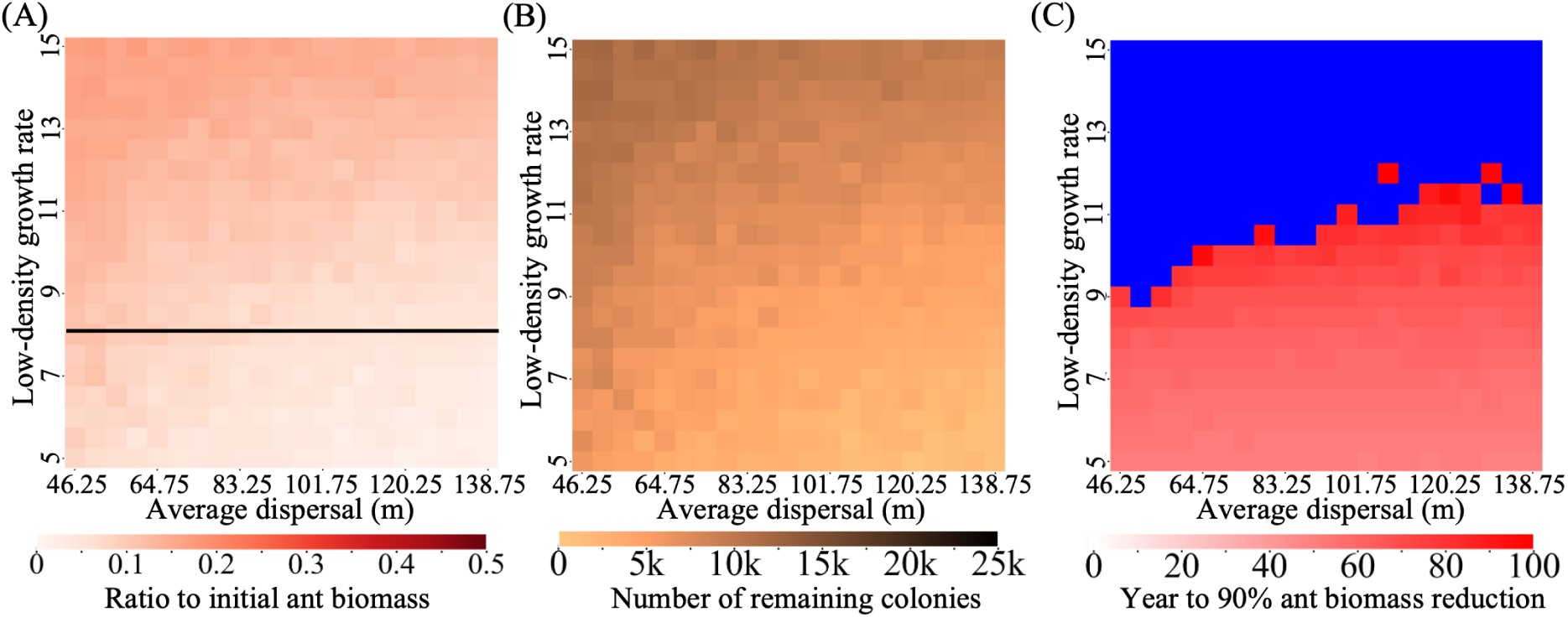
Impact of varying dispersal distance and low-density growth rate on fire ant control. The drive was released into a mixed spatial fire ant population with varying average dispersal rate (migration distance) and low-density growth rate. Heatmaps show the (**A**) fraction of remaining fire ant biomass compared to the initial biomass after 100 years, (**B**) the number of colonies remaining in the arena 100 years after the release of drive males (the initial number is 80,941), and (**C**) time required to achieve a 90% reduction in fire ant biomass. Blue indicates that the population fails to reach a 90% reduction within 100 years after drive release. The black line in panel A (and area below the line) shows where the drive has sufficient theoretical genetic load value at equilibrium to eliminate a panmictic deterministic population. Each point shows the average result from 20 simulations (including only simulations that reached 90% biomass reduction).

In contrast, a reduced low-density growth rate contributed to more effective suppression, particularly given that our drive with default parameters lacked the power to theoretically suppress populations with low-density growth rates much higher than 8. Nonetheless, all simulations within the given parameter range consistently achieved at least a 70% reduction in total ant biomass (Figure 3A), demonstrating that the default drive’s performance is highly effective in substantially suppressing fire ant populations, even with possible variations in fire ant ecology.

In polygyne colonies, newly emerged alates typically integrate into existing colonies through ambulatory dispersal rather than establishing independent colonies. This social structure allows for dynamic reproductive queen replacement within the polygyne social form. Our default values for polygyne related parameters in the model, including the representative queen replacement rate of the relative dispersal ability, were based on approximate estimates. We thus varied each of these to see how it would impact model outcomes. While polygyne queen dispersal distance minimally impacted population suppression (Figure S4A-C), it would likely have slowed down the drive wave of advance if the drive release wasn’t widespread. Our findings revealed a more nuanced relationship between queen replacement probability and population dynamics. At 30% yearly representative queen replacement probability, population suppression appears most effective (Figure S4D-F), perhaps because the drive in polygyne won’t far outstrip monogyne (due to increased generation times for polygyne), reducing chasing dynamics near the end of the simulation. However, when queen replacement probability is lower, monogyne is now reduced more quickly due to shorter generation time, though this range is unlikely to be realistic.

### Chasing in fire ants

In our previous simulations, we calculated the necessary genetic load threshold (black line) to eliminate a deterministic population (Figure 3A, 4A). Population elimination still did not occur in these simulations, so chasing is one possible reason for this. However, with the longer time scale of fire ant suppression, the effect of chasing may be less clear since it takes many generations to fully develop. To better assess this, we first examined the performance of our default drive in a panmictic model compared to our spatial model (Figure S5).

Although initial population reduction rates parallel those observed in the panmictic model, continuous space allows for higher long-term population size.

We thus extended the time scale of our spatial model to examine chasing in more detail. We found that for default parameters (where genetic load is sufficient for deterministic population elimination), the total biomass does indeed fluctuate, characteristic of chasing (Figure 6A), though population elimination did occur in a few replicates. Further, the average nearest neighbor index also fluctuated (Figure 6B), which shows varying patchiness in the population distribution and is strongly correlated with chasing^73^. Visual inspection of fire ant simulation movies also showed chasing behavior over long time scales (https://rb.gy/wnt7fn). Thus, chasing is prevalent in our fire ant models, even though it is not likely to have as large of an impact as in other models where more generations elapse in a shorter time.

**Figure 6.**
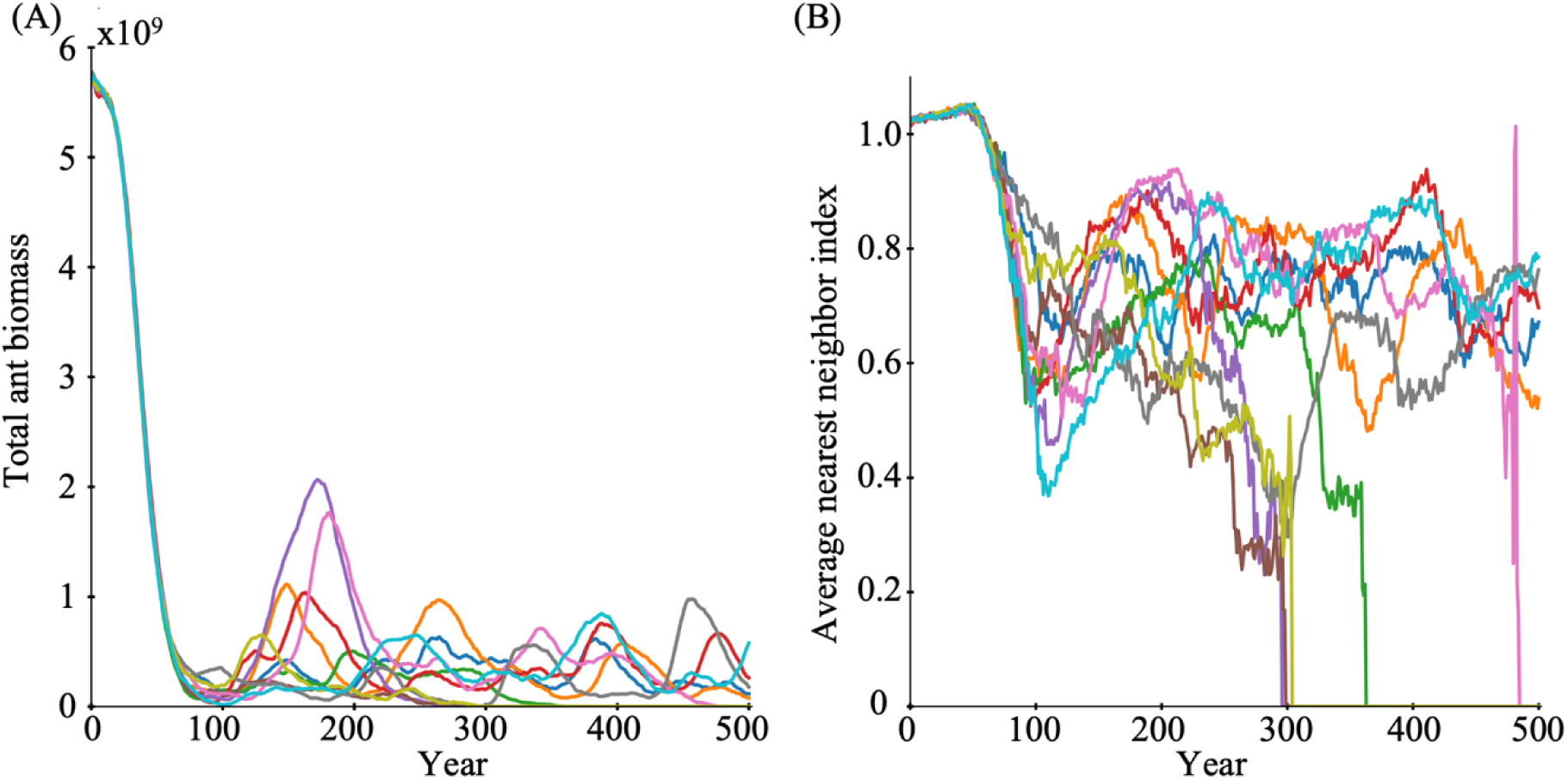
The chasing phenomenon in fire ant populations. Drive males were released into a spatial fire ant population. All model parameters were at their defaults, but the simulation was allowed to progress 500 years after the drive release. The average nearest neighbor ratio (low values < ∼0.9 represent randomly distributed individuals, which is indicative of chasing if it remains low for several generations without population elimination) and total ant biomass were tracked in each year after releasing drive males. 10 independent simulations were conducted, with each line representing a distinct simulation trajectory.

### Suppression drive designs with higher genetic load

Given the challenges in constructing standard homing drives with high efficiency by targeting recessive female-specific fertility genes, we examine alternative approaches to enhance suppression efficiency. Dominant-sterile homing suppression drives that cause dominant female-sterile resistance has demonstrated higher efficiency for self-limiting systems in modeling and in *Drosophila*^42^, but these have not yet been evaluated in haplodiploid species or in self-sustaining drives when the drive remains recessive sterile, which is highly plausible at such target sites. Here, we evaluate the genetic load of such systems, a characteristic used to measure suppressive power on a target population. In a simple discrete-generation haplodiploid model, we see that for a standard homing suppression drive (with recessive sterile resistance), high genetic load is only obtained when drive conversion is high (Figure 7A). However, for the dominant-sterile resistance version, high genetic load could be achieved even with low drive conversion, as long as the total cut rate (drive conversion + germline resistance) remains high (Figure 7B). This result is similar to two-target suppression drives in diploids^43^.

**Figure 7.**
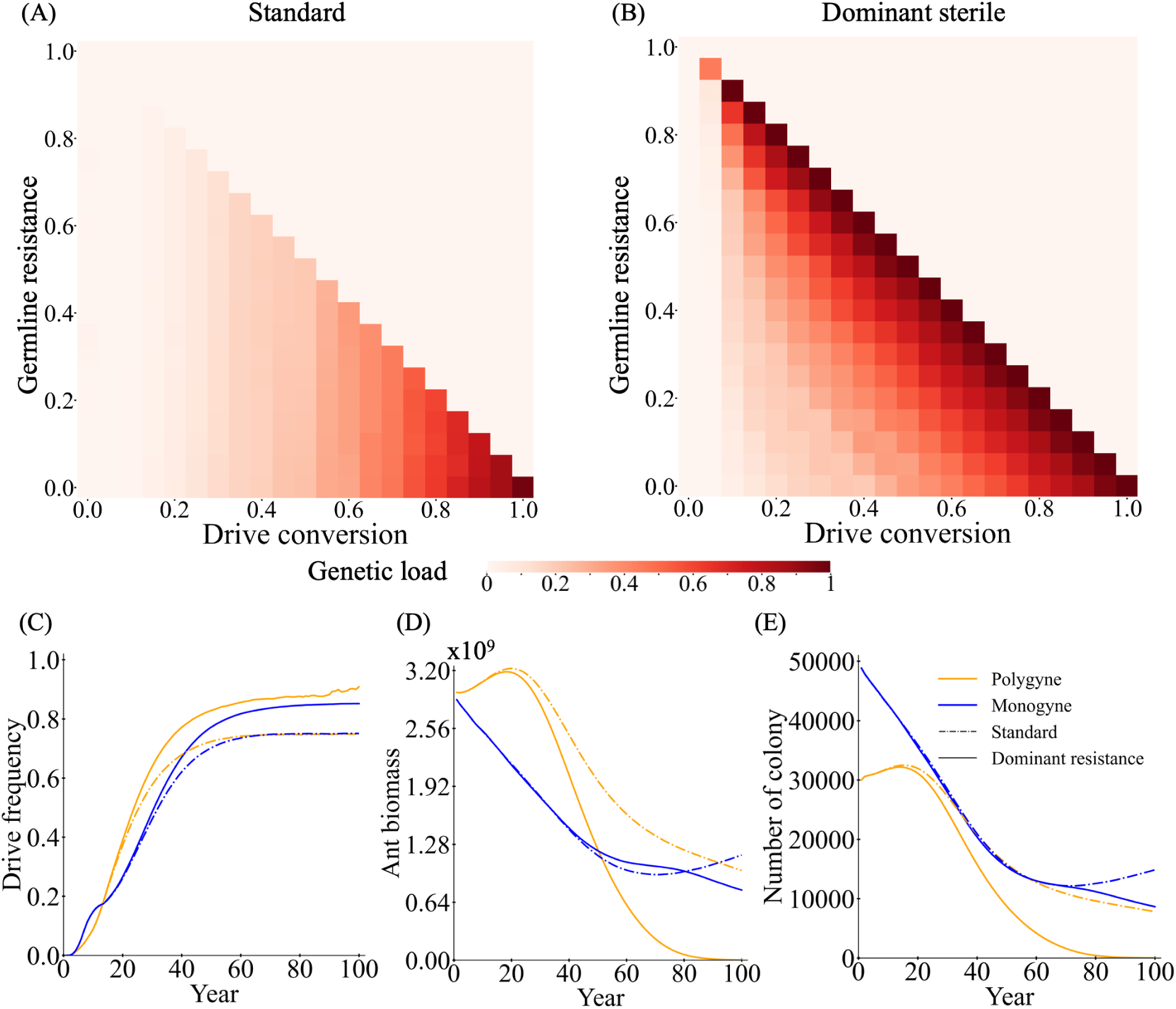
Performance of drives with dominant sterile resistance alleles. Using a generic discrete-generation panmictic model, the equilibrium genetic load (suppressive power) is shown with default performance characteristics under varying drive conversion rate and germline resistance for (**A**) standard haplodiploid suppression drives targeting female fertility and (**B**) drives that produce dominant-sterile resistance alleles. 50 simulations were assessed for each point in the parameter range. In the fire ant model with a spatial population (composed of a mix of monogyne and polygyne colonies) and a drive conversion rate of 0.8, we track the (**C**) drive frequency, (**D**) ant biomass, and (**E)** the number of colonies for each social form in simulations with standard and dominant-sterile resistance suppression drives. Displayed data are the average of 200 simulations.

While dominant-sterile resistance yields higher genetic load in the long term, the drive conversion is still the main factor in how rapidly the drive frequency increases at first, and time to population elimination is a critical consideration in fire ants. In species-specific modeling for fire ants with a less efficient drive, we nonetheless found that the dominant female-sterile resistance system demonstrates superior suppression efficacy compared to the standard drive within a few decades (Figure 7C-E), with this difference particularly pronounced in polygyne versus monogyne social forms. The dominant female-sterile resistance system exhibits comparable release ratio-dependent dynamics to those observed in standard homing drive. Increased release ratios expedite suppression across both default drives and even drives with higher fitness costs (Figure S6).

A distant-site/two-target homing suppression drive design^43^ demonstrates similarly improved genetic load based on total cut rate in haplodiploids (Figure S7). This system involves a homing rescue drive (usually considered for population modification) that has additional gRNAs targeting a female fertility gene (the same type as found in a standard homing suppression drive and specifically not a site that produces dominant-sterile resistance alleles). The two-target site design has two possible configurations, which represent a tradeoff. If the drive site gene is essential but haplosufficient, then 100% cutting will reduce genetic load, with an optimum at a somewhat lower level of total cutting (Figure S7C). Under high cutting rate conditions, the accumulation of nonfunctional resistance alleles at the drive site attenuates drive establishment, thereby slowing down the disruption of the female fertility gene and consequently reducing genetic load. If the drive site is at a haplolethal gene, which is a more difficult target for rescue, then genetic load continues to increase as the total cut rate increases (Figure S7D), comparable to dominant female-sterile resistance. Despite equivalent equilibrium genetic load at 100% cutting, the haplolethal distant-site drive system has somewhat higher genetic load when performance is less ideal (Figure S7). Further, population suppression compared to dominant female-sterile resistance is somewhat faster (Figure S8).

### Strategies to enhance suppression effectiveness by manipulating social form

Building on previous results, which demonstrate that the polygyne social form is more amenable to control, we propose that converting the social structure of fire ants could represent a promising strategy for enhancing suppression efforts. Actual implementation of such methods would require a more detailed understanding of the molecular mechanism of social form, but such investigations could be readily conducted if economical genetic modification techniques are developed for fire ants in the future.

Three strategies were designed to enhance the control of fire ants. One strategy involves converting all monogyne colonies into polygyne social form. The drive contains a dominant element that ensures that drive carries will have a polygyne colony form, even if they lack an Sb allele (Figure 8A). Sb carriers with a drive remain viable if they still have a SB allele. This approach would shift the population towards the polygyne social form, leading to a significant reduction in the number of monogyne colonies. Another approach involves a drive engineered to counteract the “greenbeard” effect by rescuing SB homozygotes in polygyne colonies if they carry a drive, thereby increasing the number of drive offspring in polygyne colonies that can lead to more rapid drive increase in the monogyne population (Figure 8B). A final approach involves targeting the SB or Sb allele for cleavage by the drive allele, promoting conversion to the alternate Sb or SB allele in germline cells that have a drive and are also SB/Sb heterozygotes (Figure 8C). When converting SB to Sb alleles, polygyne colonies may have more offspring if the queen is mated to an SB male, but if mated to an Sb male, the queen will not be viable because her offspring will all be Sb homozygotes. While monogyne colonies tend to produce more males (and wild-type polygyne colonies produce half SB males), this drive will be at a disadvantage when it reaches high frequency due to increase fractions of Sb alleles in the population. Alternatively, converting Sb to SB alleles will prevent queens from being viable is mated to an SB male, but this approach may still be better because it allows for the drive frequency of monogyne colonies to increase more rapidly, and monogyne ants are often more of a limiting factor in population suppression.

**Figure 8.**
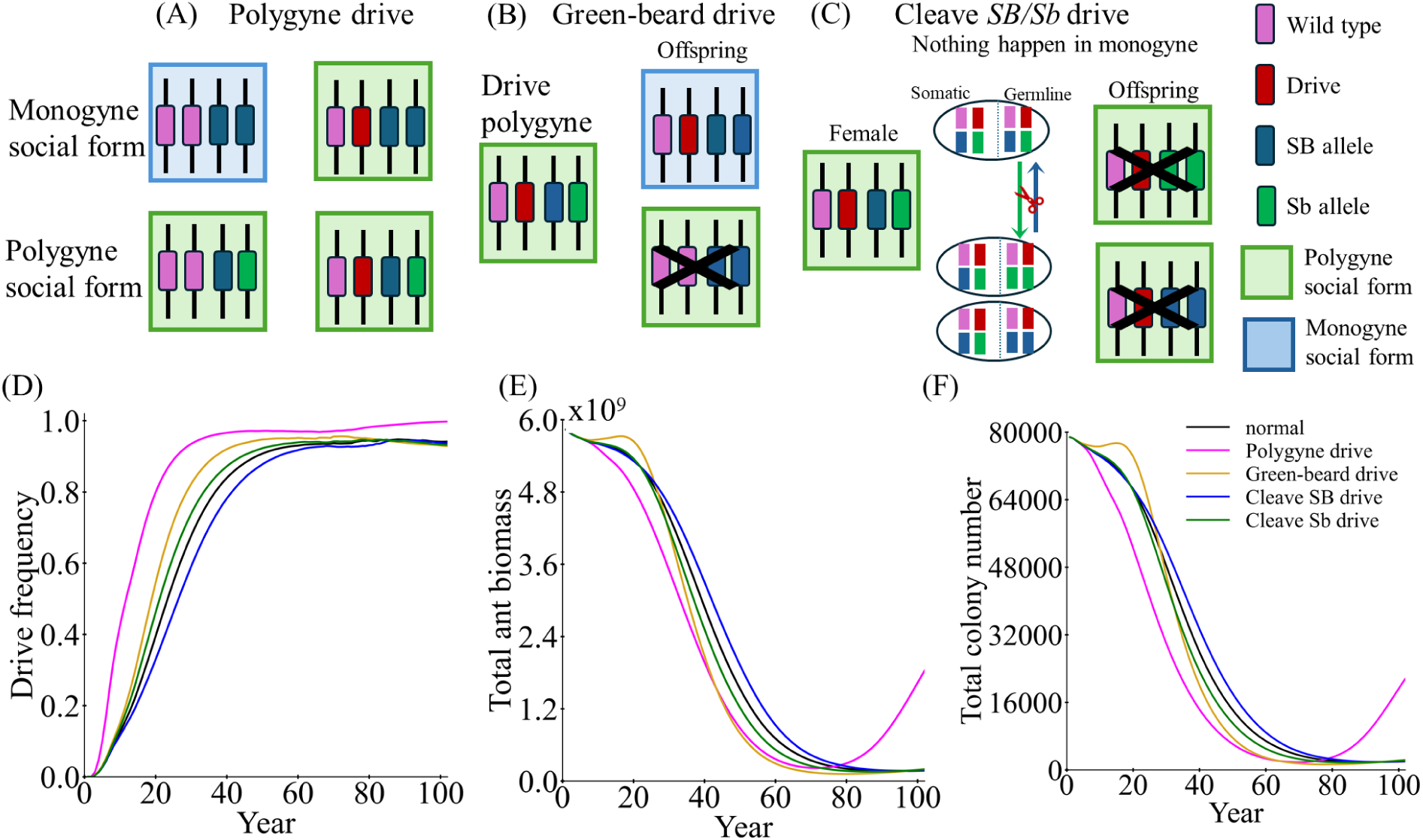
Comparison of new drive variants. (**A**) In polygyne drive, all drive queens will be polygyne, even if their genotype is *SB*/*SB*. Queens are still viable if they have at least one *SB* allele at the native polygyne locus. (**B**) In greenbeard drive, the *Sb*/*Sb* new queens born in polygyne colonies will remain viable if they have a copy of the drive. (**C**) In cleave *SB/Sb* drive, the *SB* allele or *Sb* allele is cut in the germline of drive carrying *SB*/*Sb* heterozygotes, converting it to a *Sb* or *SB* allele, which then is inherited by all offspring. Each drive was released into a mixed spatial fire ant population, and the (**D**) drive frequency, (**E**) ant biomass, and (**F**) number of colonies were tracked. Displayed data are the average of 200 simulations.

Modeling these results, we found that the polygyne drive results in a population resurgence around the 70-year mark for both social forms (Figure 8D-F, Figure S9). Though this drive increases most quickly at first, it lacks the ability to persist in monogyne, which eventually can recolonize empty space. The greenbeard mechanism shows an initial minor increase in monogyne colonies, but then demonstrates an enhanced suppression rate compared to the normal drive system, especially for the polygyne form (Figure 8D-F, Figure S9). Comparative analysis of the cleave SB drive reveals that this construct has slightly inferior efficiency relative to a standard suppression drive, but the version cleaving Sb offers somewhat superior performance (Figure 8D-F, Figure S9).

### Competing species in the fire ant model

Our previous research revealed that competing mosquito species could aid a gene drive, allowing stronger suppression and avoiding chasing^40^. As a highly invasive species, understanding the mechanisms by which fire ants invade and establish colonies in new regions, as well as how native ant populations respond to their presence, is crucial. With ants present in high numbers in almost all biomes, any invasion of fire ants is likely to be at least somewhat opposed by the native ant species, some of which may persist in a reduced ecological niche even if the fire ant invasion is highly successful. To examine the impact of such native ants on gene drive outcomes, we simulated scenarios where fire ants and native ants coexist (though with a higher starting fire ant population), with symmetrical competition factors, representing the relative competition strength between the two species. We found that the presence of native ants contributes substantially to the suppression of the fire ant population, facilitating successful suppression with greater competition (Figure 9A-B). In fact, complete fire ant elimination becomes possible, though this still requires a temporal window exceeding 80 years (Figure 9C). This timeline suggests the necessity for long-term management strategies despite the beneficial effects of native ant competitors. On the other hand, it also suggested that good ecosystem management maximizing the health of native species may allow greater success in gene drive deployments against invasives.

**Figure 9.**
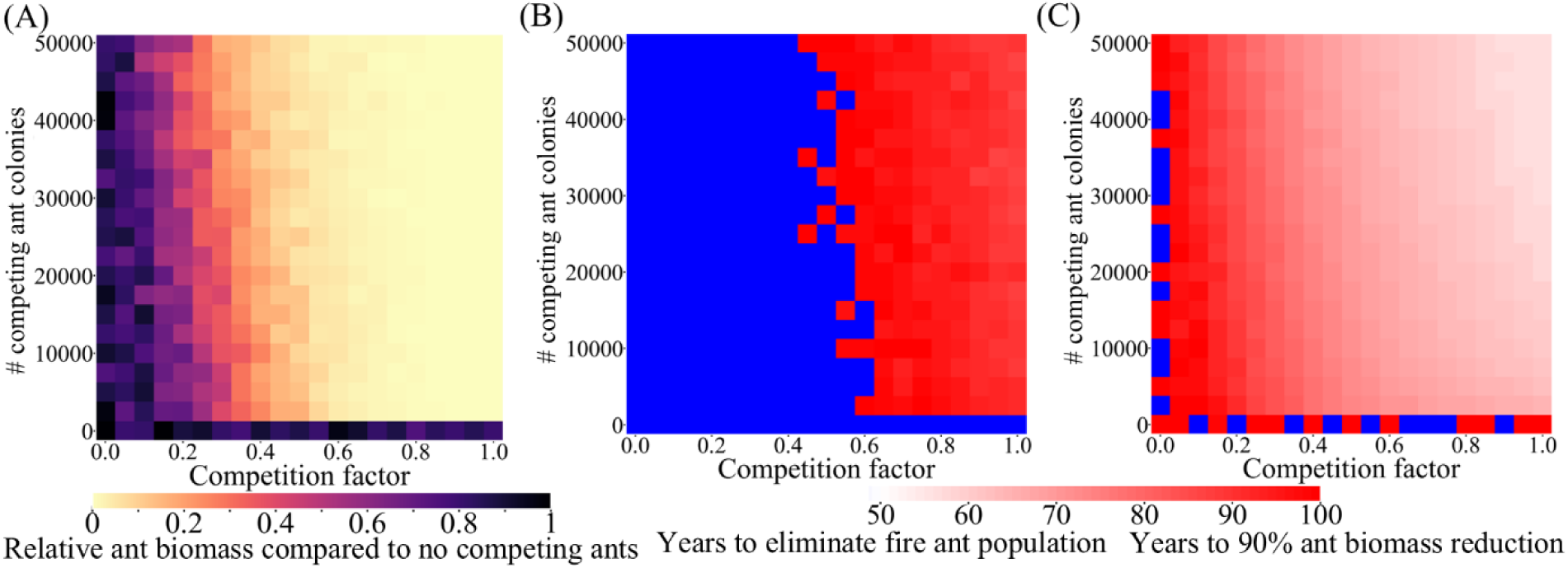
Gene drive suppression of fire ants is boosted by a competing species. The drive with default parameters (except for the low-density growth rate, which was increased to 10) was released into a spatial mixed fire ant population. The interspecific competition factor and equilibrium number of competing ant colonies was varied. Heatmaps show (**A**) the relative fire ant biomass at year 100 after drive release (compared to identical simulations without competing ants), (**B**) the time to population elimination following drive release (usually close to 100 years), and (**C**) the time to achieve a 90% reduction in fire ant biomass, with persistence beyond 100 years indicated in blue. Displayed data are the average of 20 simulations (including only simulations with population elimination).

### Drive deployment to halt a fire ant invasion

In some cases, native ants may not have a unique ecological niche to allow them to partially persist in the face of a fire ant invasion. They will then tend to only coexist with fire ants for limited time scales on the leading edge of an invasion. Here, we started a scenario where such weaker native ants are being pushed back by fire ants, with slower dispersing polygyne fire ants following behind monogyne (Figure 10A). Drive release was initiated when fire ant populations had partially advanced into the native population after the start of the simulation, allowing us to evaluate the efficacy of gene drive intervention. Monogyne colonies were the primary contributors to the high invasive strength of fire ants due to their rapid territorial expansion through nuptial flights.

**Figure 10.**
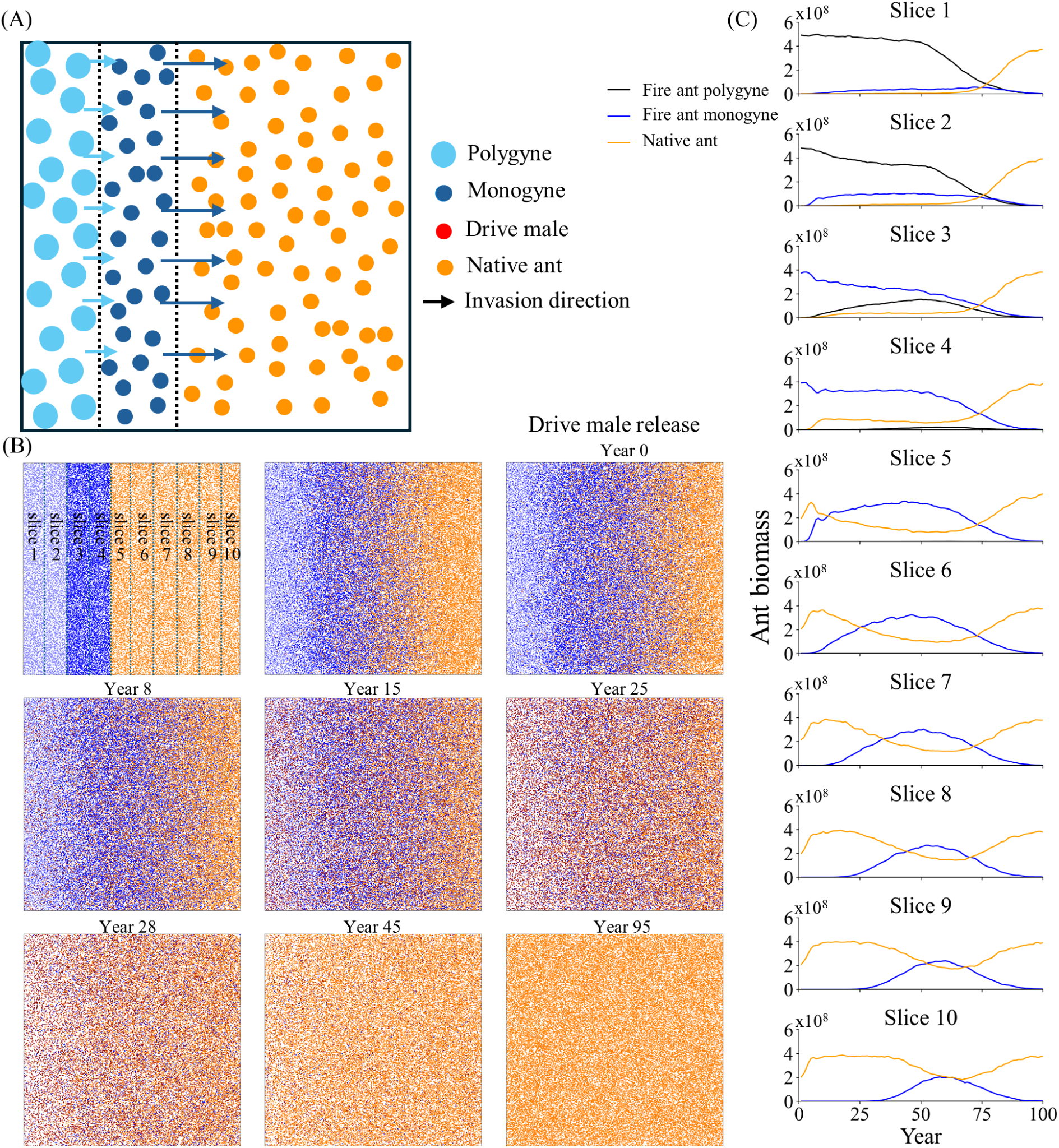
Gene drive against an invading fire ant population. (**A**) A diagram of the simulation start and invasion process. Red gene drive individuals are released in the population when the biomass in the left half of slice 7 is composed of 50% fire ants. The arrows indicate the invasion speed, which is proportional to the average dispersal. Native ants suffer full competition from fire ants, but their competition on fire ants is reduced by half. (**B**) Snapshots of the invasion process. The blue dashed lines indicate 10 slices used for tracking ant biomass. (**C**) The ant biomass of each 10% slice for native ants and each social form of fire ants. Year 0 is the year of drive release.

They swiftly advanced, but eventually lost their competitive advantage when the drive reached high frequency, retreating and rapidly declining in frequency (Figure 10B). In contrast, polygyne colonies exhibited more limited dispersal and were also eliminated more quickly by the drive. Through the implementation of a suppression gene drive, native ant populations were able to reclaim the entire territory, even in regions that had previously been completely colonized by fire ants, though the drive needed to progress for a few decades before native ants began to recover (Figure 10C).

### Identification of female fertility target genes in fire ants

Due to the difficulty in generating constructs in fire ants, even assuming steady future advances in genetic engineering technology, it will be important to use related gene drive studies to more rapidly generate viable gene drives. We thus considered several female fertility target genes in fire ants as potential candidates, desiring highly conserved regions, sites for implementing a multiplexed gRNA strategy, and previous demonstrations of suppression drives targeting homologs other insects^74^. Potential target genes included the female-specific exon of *doublesex*, *intersex*, *virilizer*, *Octopamine β2 receptor*, *NADPH oxidase*, *stall*, and *nudel*.

Multiple sequence alignment was performed across species including *Solenopsis invicta*, *Linepithema humile* (invasive Argentine ants), *Aedes aegypti*, *Anopheles stephensi*, *Drosophila melanogaster*, *Vespula vulgaris* (invasive wasps), *Bemisia tabaci* (the whitefly, an agricultural pest), and *Spodoptera frugiperda* (the fall armyworm, also an agricultural pest).

Two of these genes, *stall* and *nudel*, failed to exhibit substantial regions of conserved sequence (Figure S10). The female-specific exon of *doublesex* also exhibited poor nucleotide-level conservation, in contrast to its high conservation between many other species^24^. Therefore, only the fire ant ortholog of this sex-specific exon was directly employed for gRNA design, targeting early in the exon to potentially generate dominant-sterile resistance alleles^42^. Highly conserved regions of the remaining four genes were selected for the design of gRNA target sequences. Such regions that are strongly associated with evolutionary conservation across species tend to have high functional significance.

Thus, targeting them likely favors the formation of nonfunctional resistance alleles instead of functional resistance. Nucleotide sequences of conserved regions in fire ants were subsequently identified and used for guide RNA target site selection and design (Figure 11). These were designed for maximum multiplexing efficiency, placing target sites as close together as possible while still keeping to a low level the chance that a mutation at one gRNA target site will block another gRNA from binding.

**Figure 11.**
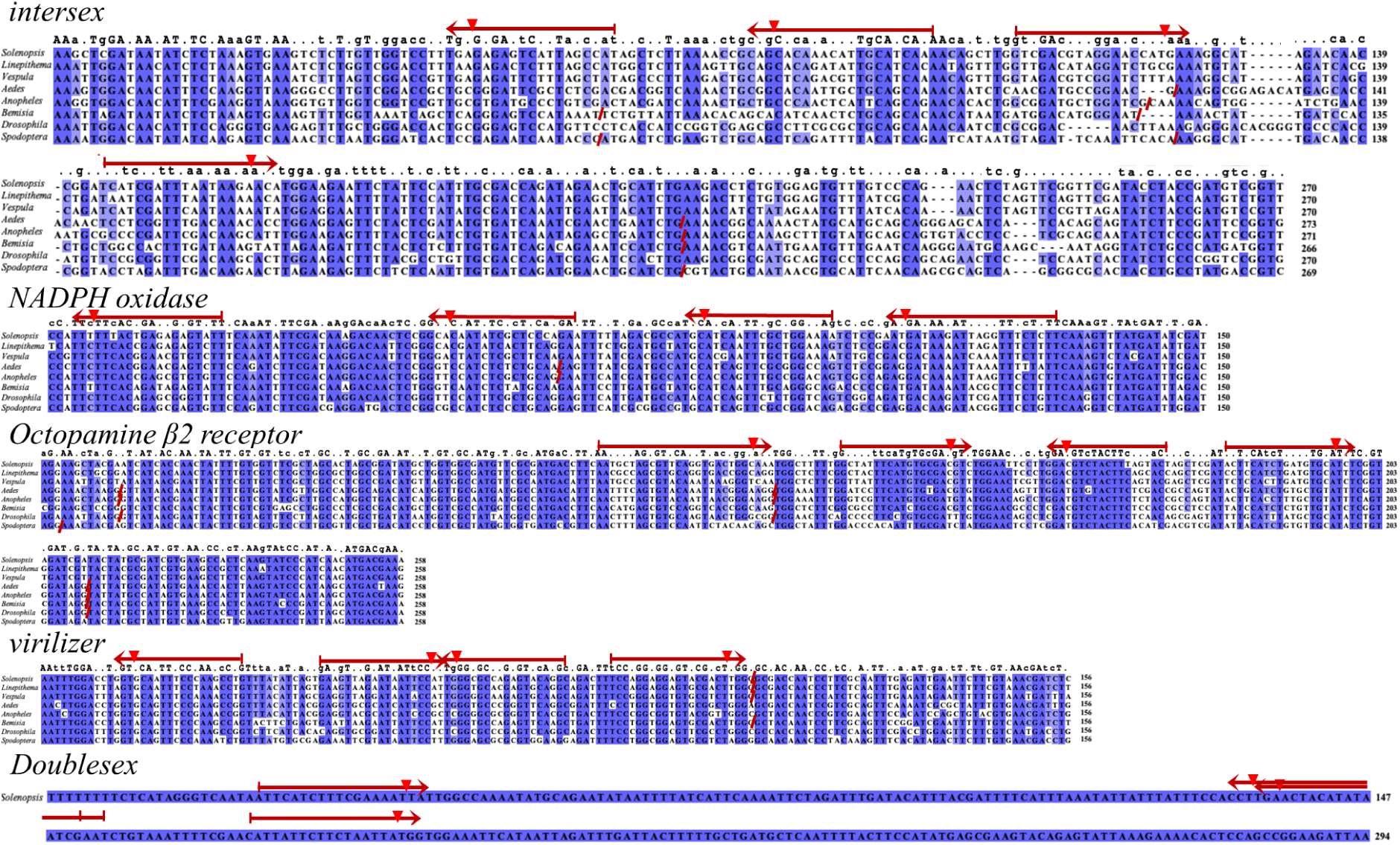
gRNA multiplexing targets in fire ant female fertility genes. Female fertility genes with highly conserved regions between several major insect species were identified. Within these regions, four fire ant target gRNA sequences suitable for a gRNA multiplexing strategy were identified using the Chopchop platform, indicated by red arrows. The gRNA cut sites were indicated by reversed triangle. The locations of introns in other species are labeled with red slashes.

## Discussion

The fire ant has garnered significant attention as an exceptionally challenging invasive species to control, characterized by remarkable ecological adaptability and robust population maintenance mechanisms. In this study, we systematically investigated the potential of suppression gene drive as a novel population management strategy for this species. Our analysis demonstrates the potential efficacy of suppression gene drive in managing fire ant populations, albeit with a substantially extended time scale compared to other gene drive applications due to the fire ant’s relatively long generation time for an insect.

The propagation dynamics of homing drive are significantly influenced by the reproductive strategies of the target species^41,75^. In fire ants, queens engage in a single mating event and retain sperm for the reminder of their reproductive lives without remating^48,76,77^. Queens live several years, and colonies tend to produce few reproductive individuals until they reach large size, thus increasing the generation time and complicating suppression. This slows suppression gene drive, but it also potentially makes it easier to achieve high release rates, since several yearly releases can take place within a single generation. Such high releases would be useful for achieving suppression on reasonable time scales, and rearing fire ants in the lab can be performed readily, but to enable releases of males during the mating season, it is possible that semi-field rearing strategies would be required so that released drive individuals perform their nuptial flight at an appropriate time. Moreover, even the release of new queens would not result in an initial increase in ant biomass, which is advantageous as it eliminates the need for sex sorting and monitoring the precise timing of mating seasons (semi-field colonies at a release site could be allowed to reproduce naturally). Matings between polygyne queens and released drive males or even direct release of polygyne queens may be particularly valuable because polygyne queens could join established nests, allowing them to produce drive alates more quickly.

Fire ant social form proportions vary significantly across invaded regions. Surveys in Florida and southeastern U.S. states reveal stable, polygyne-predominant populations^64,78^, while South America shows a monogyne-dominant pattern^77^. In China, social form proportions differ across distribution ranges^79^. Our simulations demonstrated that polygyne populations exhibited increased susceptibility to suppression, indicating that social structure composition substantially influences drive system efficacy. This dynamic is likely due to their shorter generation time of polygyne queens, which live shorter and share the nest with other queens that may be continually arriving each year^80^. In territories dominated exclusively by polygyne social forms, our model predicts complete population elimination, resulting in successful habitat restoration (Figure 2). Targeted strategies can exploit this vulnerability to facilitate suppression in mixed populations. In some drive systems designed to induce social form transformation, population suppression efficiency could be enhanced. Indeed, such greenbeard mechanisms are prevalent among eusocial insects and are often vulnerable to “false beard” cheating^81^. By exploiting this phenomenon, we tested a system that manipulates the greenbeard mechanism, allowing monogyne new queens with the drive to survive greenbeard effects in polygyne colonies, enhancing suppression efficiency by allowing polygyne colonies to more heavily contribute to drive frequency in monogyne populations, which are the limiting factor for population suppression. These drive designs may currently be difficult to implement in practice, though they may become possible, together with other novel alternatives, as more studies are conducted on the mechanism behind social form behavior in fire ants and related species. For example, it might be possible to design transgenes to allow monogyne drive colonies to escape from aggressive interactions with polygyne colonies.

To further enhance suppression outcomes, two additional improved drives (dominant-sterile resistance and two-target/distant-site drives) were developed based on the standard homing suppression drive. Although careful design is required to achieve dominant-sterile resistance^42^, it has the potential to achieve a higher genetic load compared to other drive systems with the same parameter set. The two-target design offers similar improvements but requires additional gRNAs compared to a standard drive^43^.

While our model incorporates a standard dispersal parameter, nuptial flight distances in fire ant exhibit considerable individual variation, with documented dispersal ranges demonstrating significant heterogeneity during reproductive flights^49^. However, our results demonstrated that with widespread drive release (likely necessary for drive success on acceptable time scales) the nuptial flight distance exerts minimal influence on suppression outcomes, though dispersal likely assumes greater ecological significance during invasions, and mostly likely would affect long-term chasing dynamics in fire ants too if they manage to persist in the face of the drive and other pressures.

Biological invasion represents a complex, sequential process, including transport, introduction, establishment, and spread^82^. Each stage of invasion is characterized by ecological constraints and selective pressures that critically influence the potential success or failure of an invasive species^83^. The release of gene drive during the invasion process interferes with the progression of the invasion, facilitating the elimination of fire ants and recovery of native ants. Indeed, interspecific competition may play a more substantial role in regulating fire ant population within shared ecosystems. These dynamics increase the chance of successful suppression outcomes^40^. Even if fire ants tend to be dominant, native ants can provide at least some significant competition^84,85^, and models must incorporate this process. Specifically, studies have shown that fire ant territory size is highly modulated by interspecific competitive interactions, especially the relative agonistic capacities of adjacent heterospecific colonies^86^. The presence of these native species can reduce colony establishment success in fire ants by increasing the likelihood of brood raiding during colony foundation and incipient stages^87^, just as in our model.

Despite incorporating multiple ecological factors, our model has limitations regarding niche-specific demographic and geographic variables that may influence drive suppression outcomes. Landscape heterogeneity may influence fire ant transmission dynamics and release patterns. Uncertainly in understanding interactions between social forms across different regions could also affect ultimate outcomes. We also assumed long-term persistence of polygyne colonies, but ant habitats may be more widely variable, with natural extinction-recolonization cycles that may facilitate gene drive suppression by reducing effective generation times. Many of our model parameters were also estimated based on a limited number of studies. Future research could investigate these factors in more detail, allowing better understanding of fire ant control methods while also providing fundamental knowledge about various aspects of this interesting species. Modeling could also potentially be used to investigate the mechanisms of monogyne and polygyne coexistence by adding additional features and examining parameter ranges where their population dynamics more precisely match ecological field studies.

To actually deploy gene drives in fire ants or other invasive ants such as Argentine ants or crazy ants, additional advances will be needed in ant genetic engineering. However, long-term prospects are promising in this field. Genetic knock-outs have been achieved in fire ants^88^, and knock-ins in another ant species^89^. One major limitation is rearing ants in the lab that produce alates at appropriate times. Additional research may overcome this, but more resource-intensive semi-field/outdoor conditions may be needed to ensure full lifecycles among research subjects. Such considerations also apply to actual releases of fire ants. One possible way of overcoming this may be to rear several colonies in the lab and then release them after a few years, avoiding early colony mortality and allowed each “released” colony to produce many alates over the course of a few years.

Overall, we conclude that gene drive in fire ants is promising but likely to be a highly challenging application, both due to the difficulty in generating the needed ants and releasing them at large scale. Even then, time scales for high population suppression will optimistically still extend over a few decades. Thus, other short-term fire ant control research remains essential. However, long-term outcomes for gene drive appear to be positive, particularly with improved variants, and other methods of control may be less efficient and more resource intensive, even if they yield more immediate benefits. Thus, long-term investment in genetic and rearing technique research could yield major long-term benefits for gene drive-based control in fire ants and many other major invasive ant species.

## Acknowledgements

We would like to thank Khushi Patel and Xuejiao Xu for their assistance. We also gratefully acknowledge the High-Performance Computing Platform of the Center for Life Science at Peking University for data collection. This study was supported by the Center for Life Sciences, as well as the grants from the National Science Foundation of China (32270672 and W2432018) and the SLS-Qidong Innovation Fund.

## Supplemental Information

**Figure S1.**
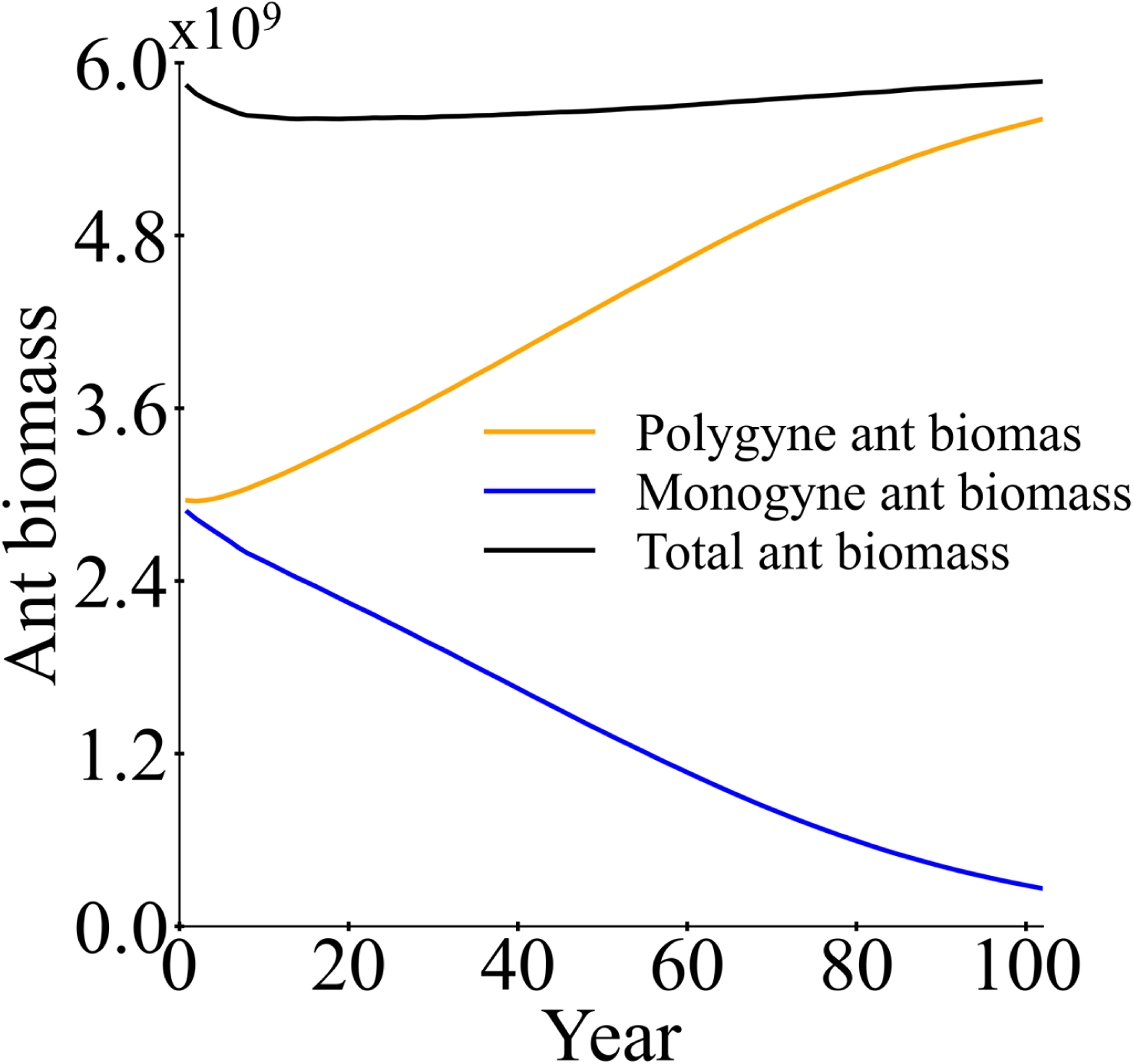
Total ant biomass without releasing drive. The total ant biomass in a mixed population (monogyne and polygyne) was recorded in each year without releasing drive individuals. One unit of biomass is the mass of one monogyne colony worker. Displayed data are the average of 20 simulations.

**Figure S2.**
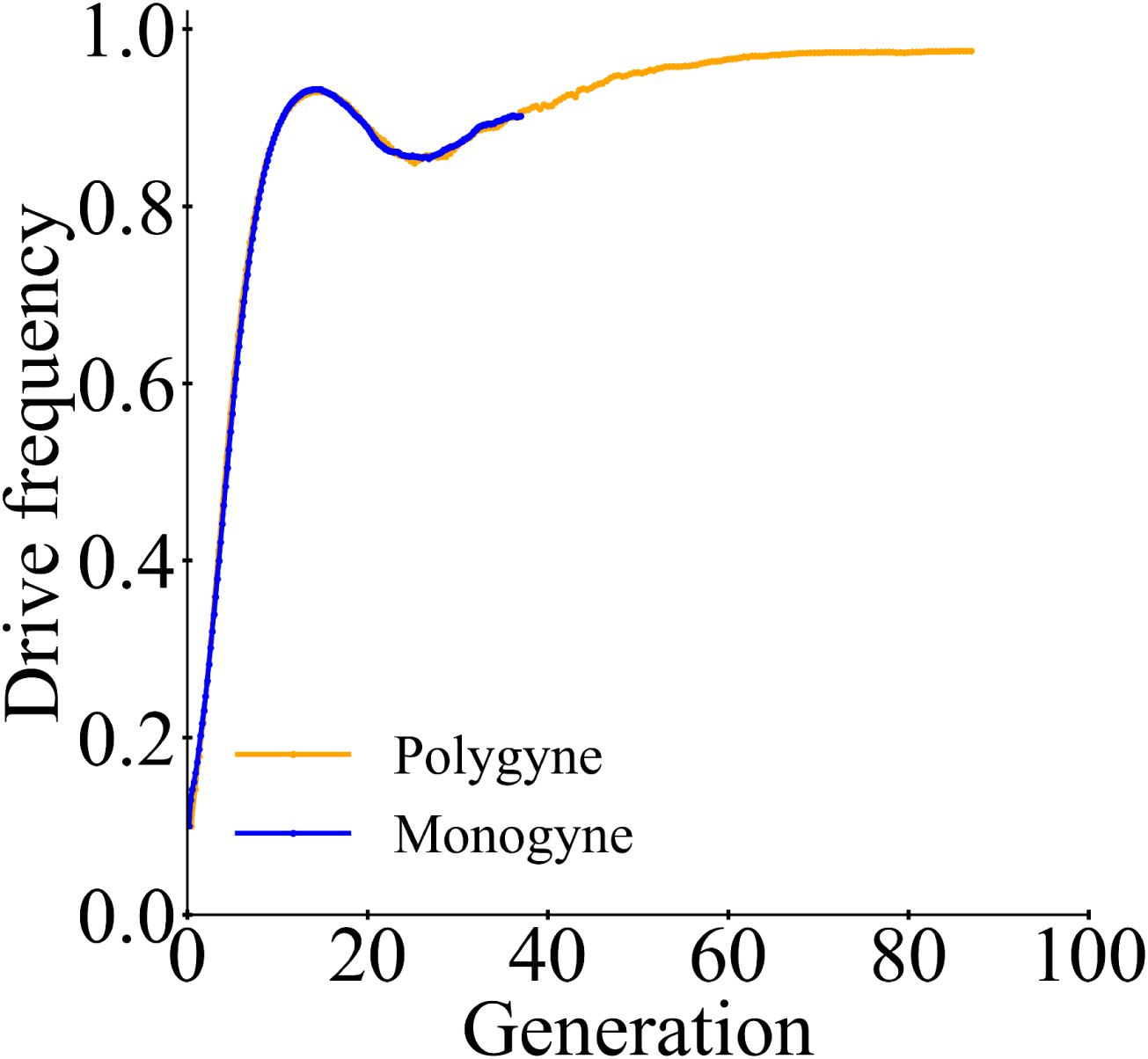
Comparison of generation time. The drive frequency in monogyne-only and polygyne-only models was tracked in each year. The initial population contained 10% drive heterozygous and 90% wild-type colonies. The simulation was run for a duration of 200 years, and these were converted to generations. For monogyne, the generation time was calculated as 5.41 years. The generation time for polygyne should be slightly higher than 2 years based on 50% queen replacement, and it was found that allele frequency would exactly match monogyne with a generation time of 2.3 years, as shown. Displayed data are the average of 200 simulations.

**Figure S3.**
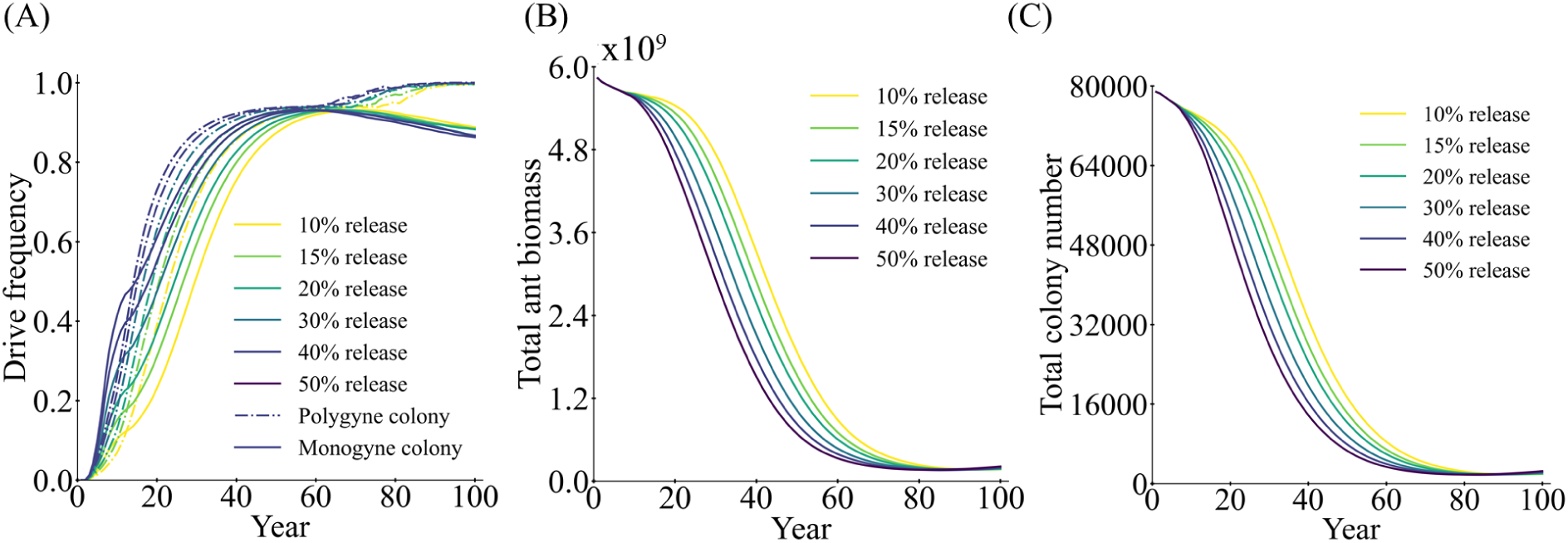
Effect of drive release size. With varying drive male release sizes over six years (as a fraction of the total population), we tracked the (**A**) drive frequency for polygyne (dashed lines) and monogyne (solid lines) social forms, (**B**) total ant biomass and (**C**) total colony number. Displayed data are the average of 200 simulations.

**Figure S4.**
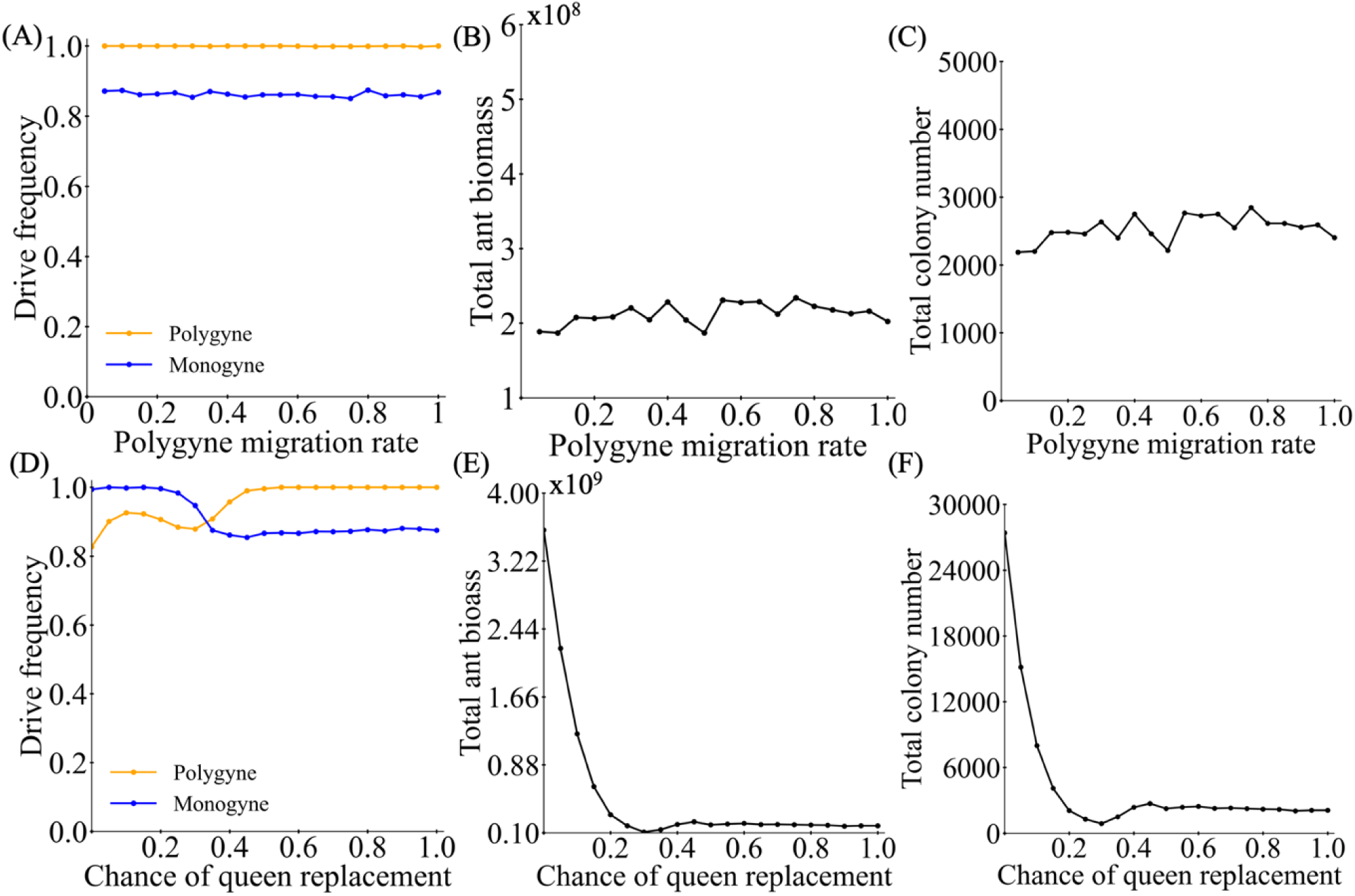
Suppression outcomes with varying polygyne colony parameters. We varied the (**A-C**) relative dispersal rate of polygyne ants compared to monogyne and varied the (**D-F**) chance of representative queen replacement in polygyne colonies. We tracked the (**A,D**) drive frequency, (**B,E**) total ant biomass, and (**C,F**) total colony number. Displayed data are the average of 200 simulations.

**Figure S5.**
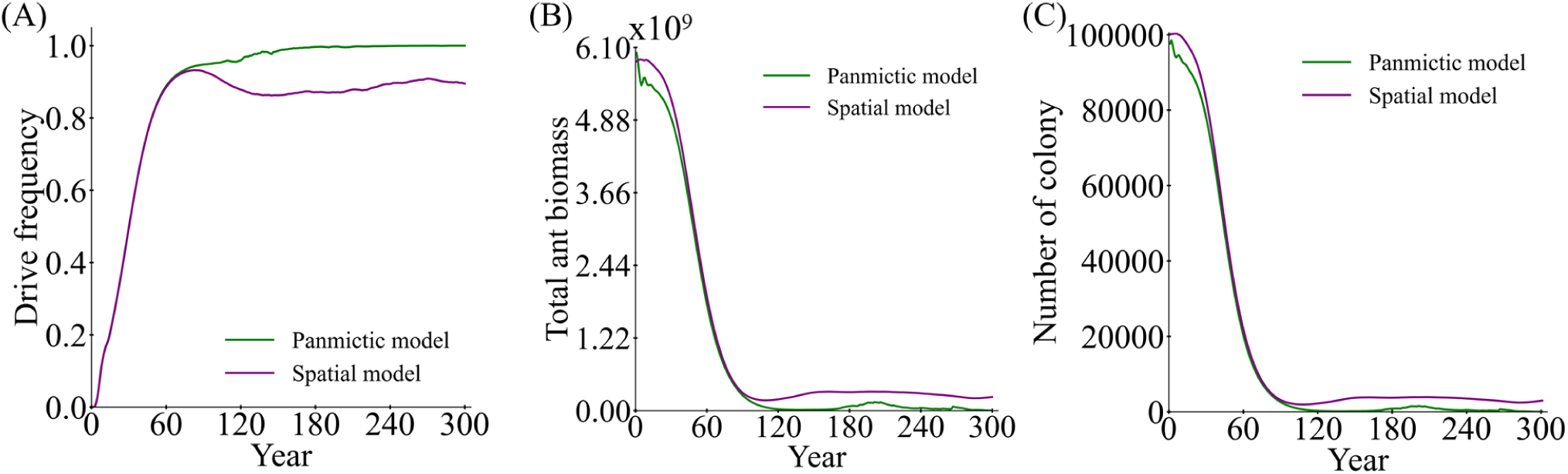
Comparison of panmictic and spatial fire ant models. Using default parameters and monogyne only populations in our normal spatial model and in a panmictic model, we tracked the (**A**) drive frequency, (**B**) total ant biomass and (**C**) number of colonies. Each point shows the average of 200 simulations.

**Figure S6.**
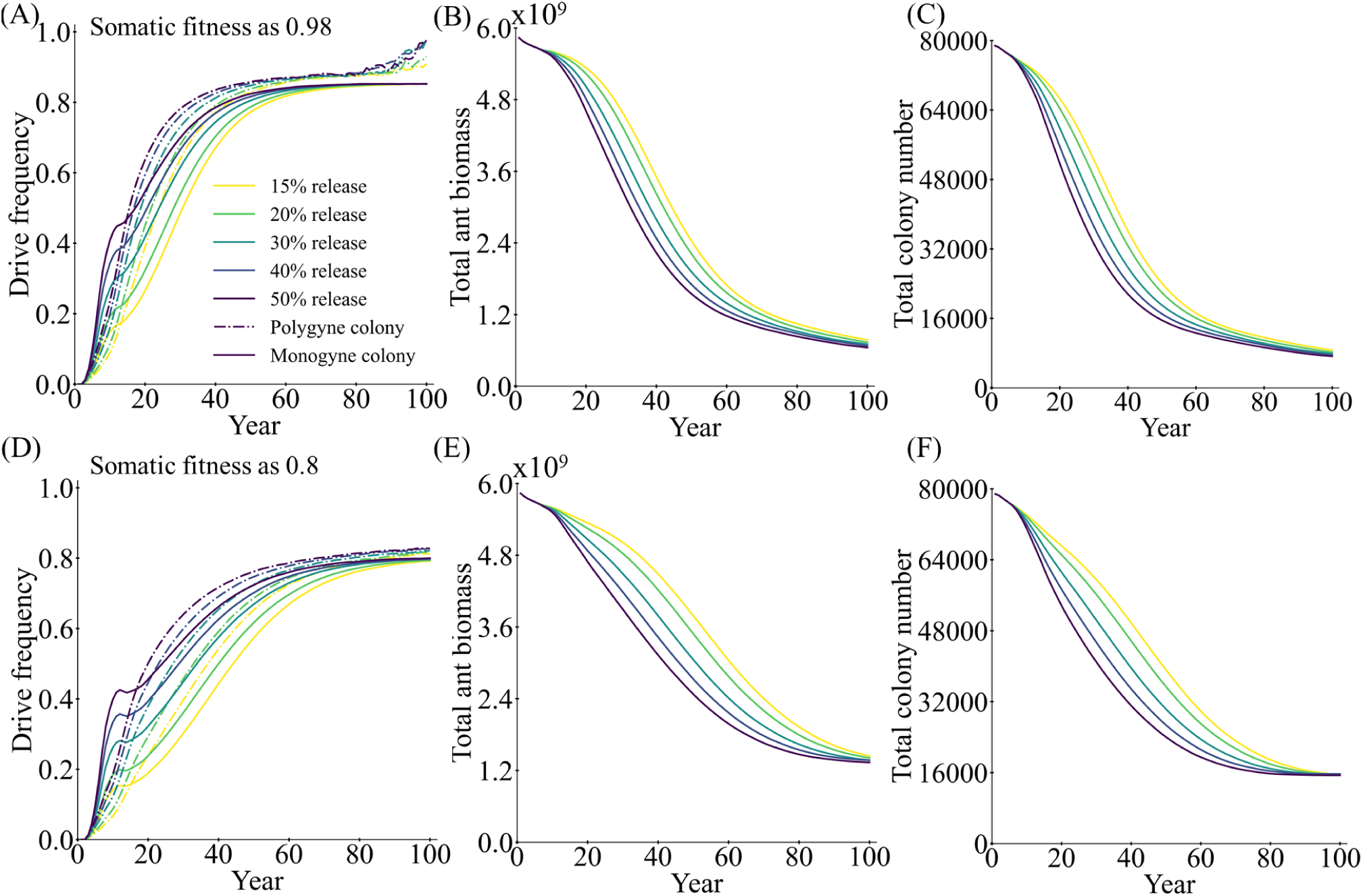
Different release ratios of dominant sterile resistance drive. In the fire ant model with a mixed spatial population (composed of a mix of monogyne and polygyne colonies), a drive conversion rate of 0.8, and a female fitness of (**A-C**) 0.98 or (**D-E**) 0.8, we release either standard or dominant-sterile resistance suppression drives. We track the (**A,D**) drive frequency, (**B,E**) ant biomass, and (**C,F)** the number of colonies for monogyne and polygyne social forms. Displayed data are the average of 200 simulations.

**Figure S7.**
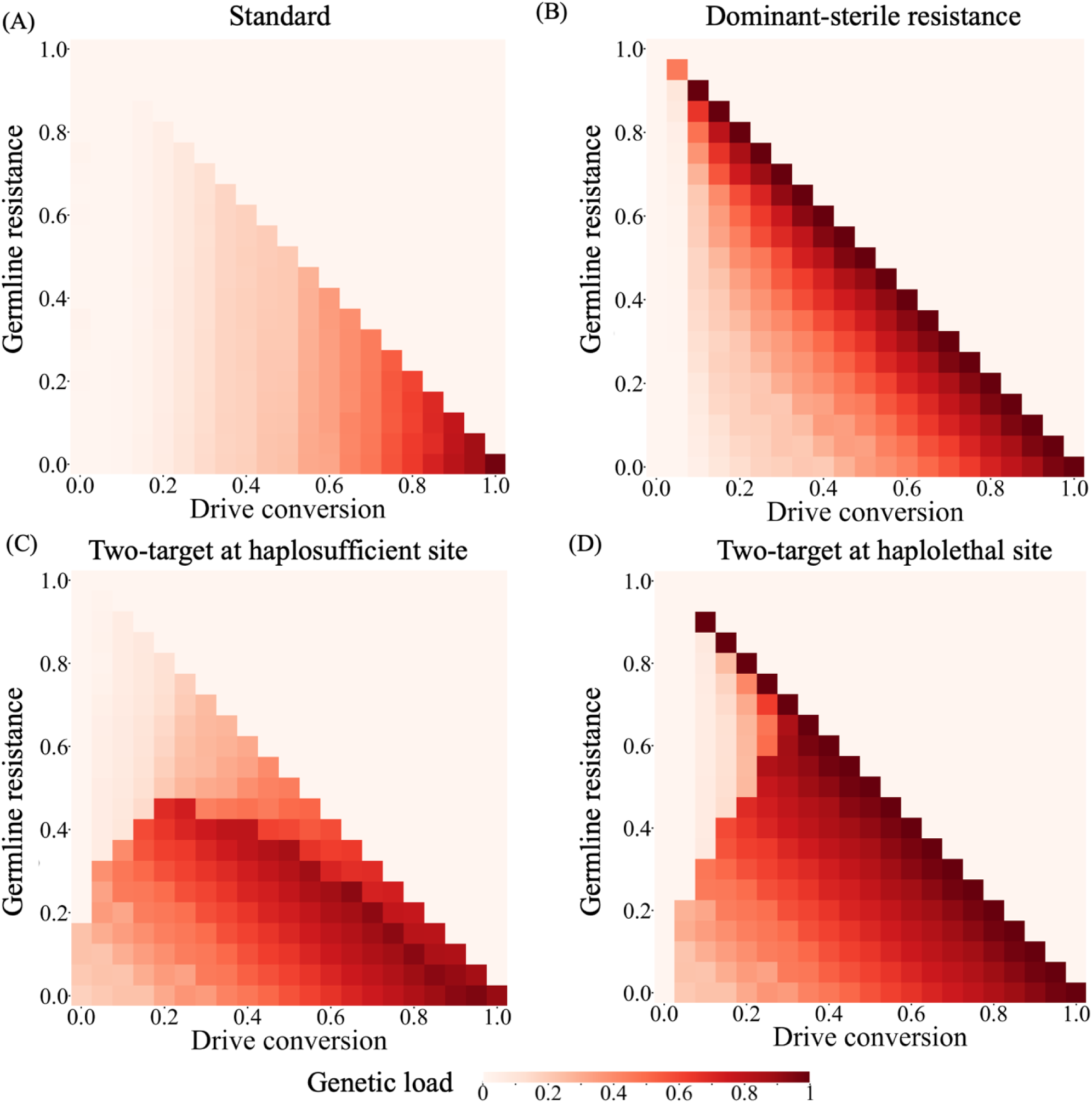
Suppression efficiency comparison of four suppression drive designs. Using a discrete-generation panmictic model, the equilibrium genetic load (suppressive power) is shown with default performance characteristics under varying drive conversion rate and germline resistance for (**A**) standard haplodiploid suppression drives targeting female fertility, (**B**) drives that produce dominant-sterile resistance alleles, (**C**) two-target design with the drive in a haplosufficient but essential gene, and (**D**) two-target design with the drive in a haplolethal gene. (A) and (B) are identical to Figure 7 to facilitate direct comparison. Data points in (C) and (D) are the average of 20 simulations.

**Figure S8.**
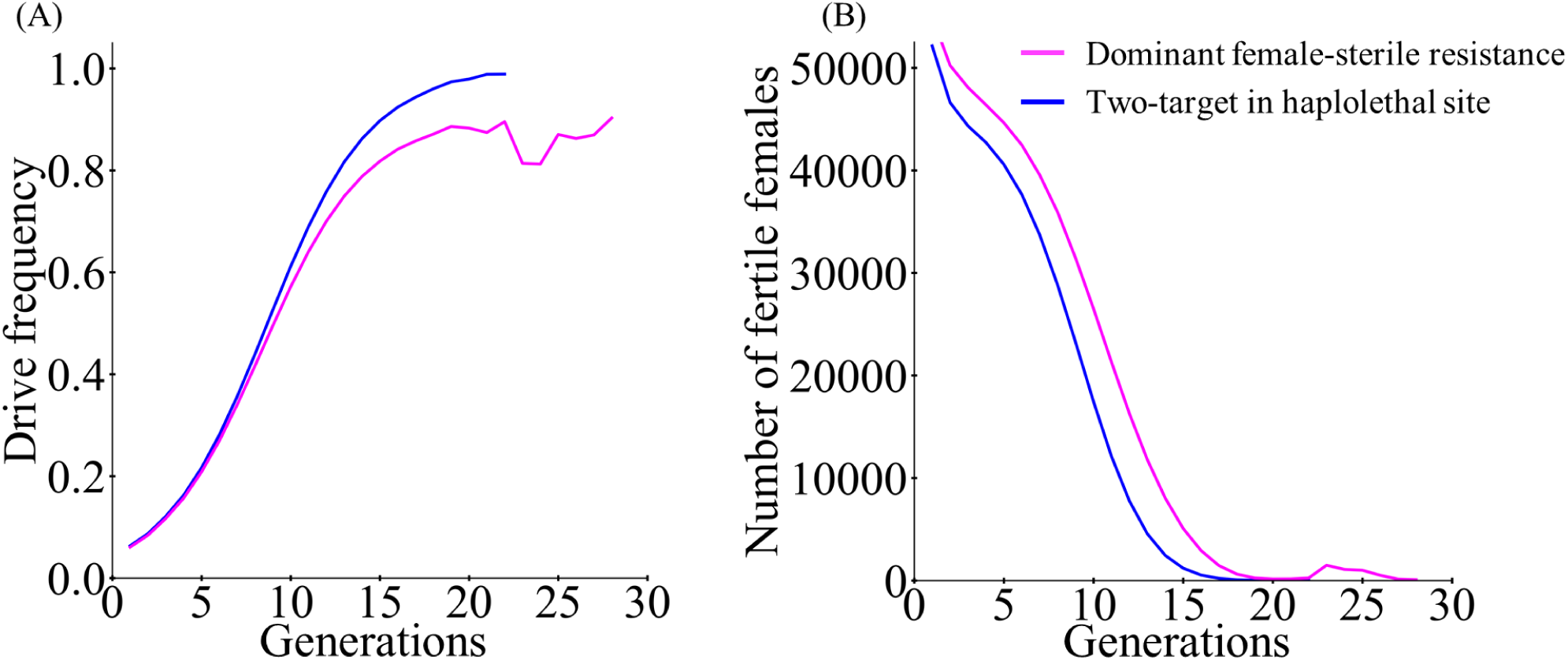
Comparison of haplolethal-suppression distant drive and dominant female-sterile resistance drive. Using the discrete-generation panmictic model, the **(A)** drive frequency and **(B)** population size in each generation are tracked with default performance characteristics, when total cut rate is 1. Displayed data are the average of 200 simulations.

**Figure S9.**
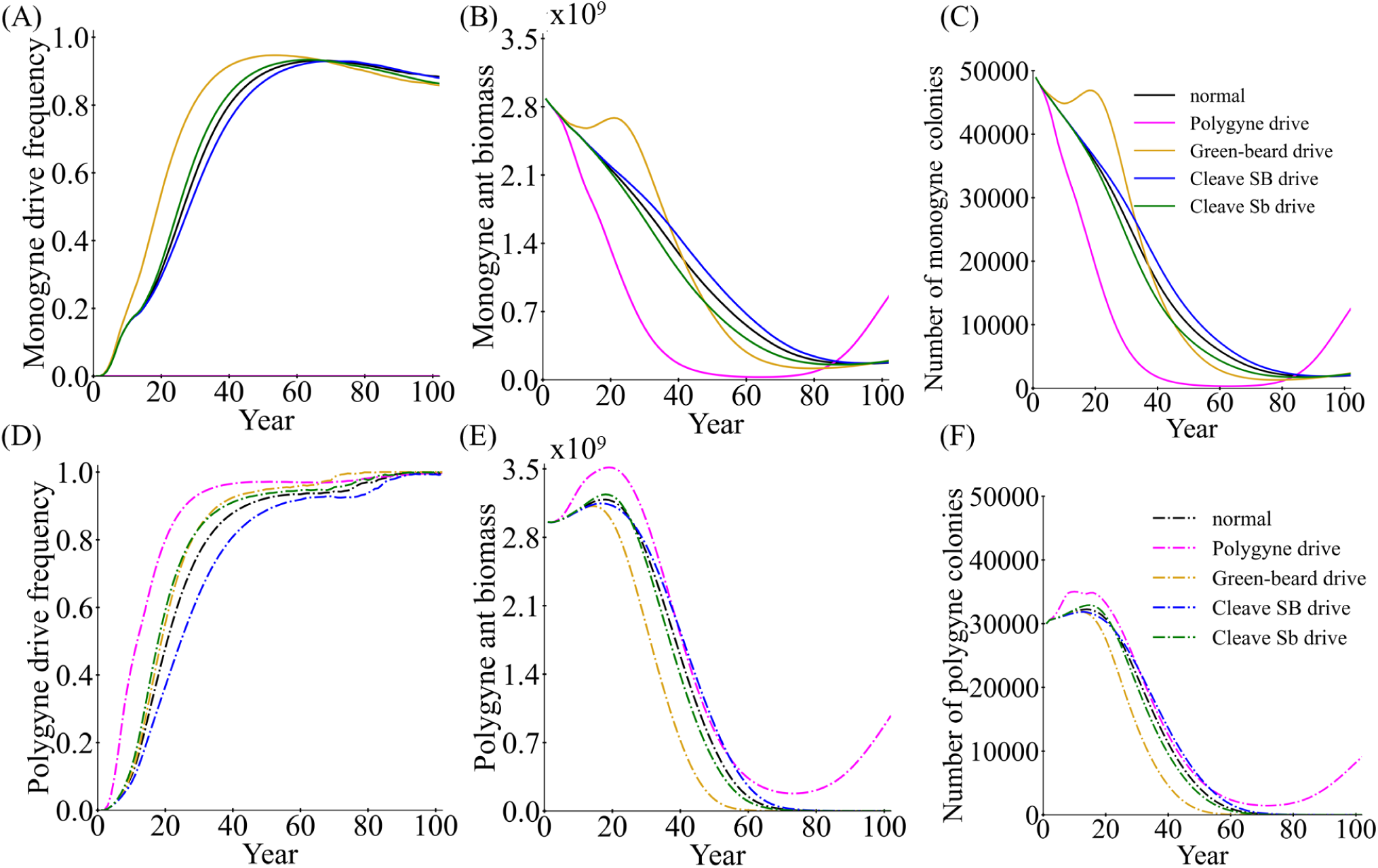
Comparison of new drive variants. Each drive variant was released into a mixed spatial fire ant population. For (**A-C**) monogyne and (**D-F**) polygyne social forms, we track the (**A,D**) drive frequency, (**B,E**) ant biomass, and (**C,F**) number of colonies. Displayed data are the average of 200 simulations.

**Figure S10.**
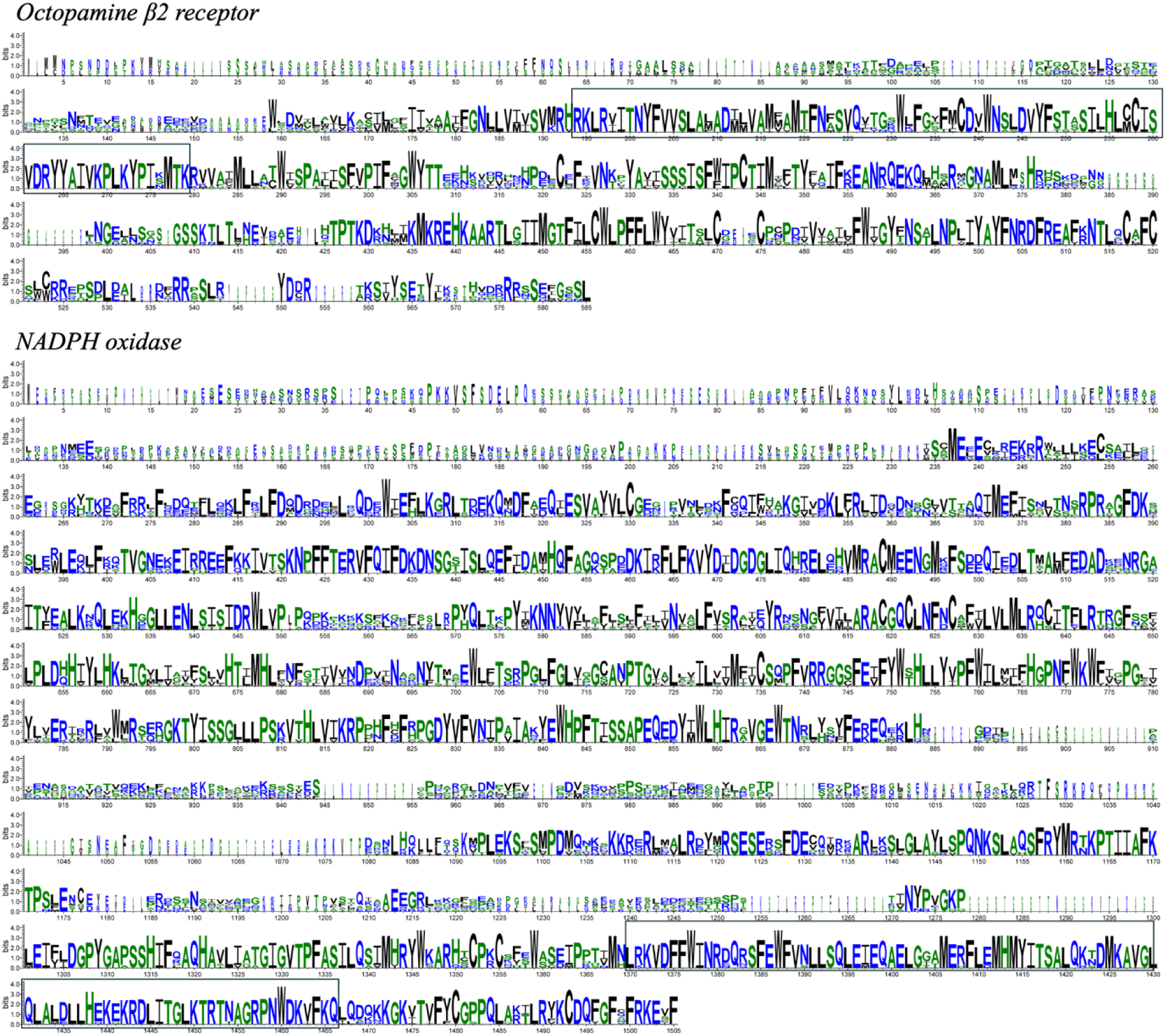
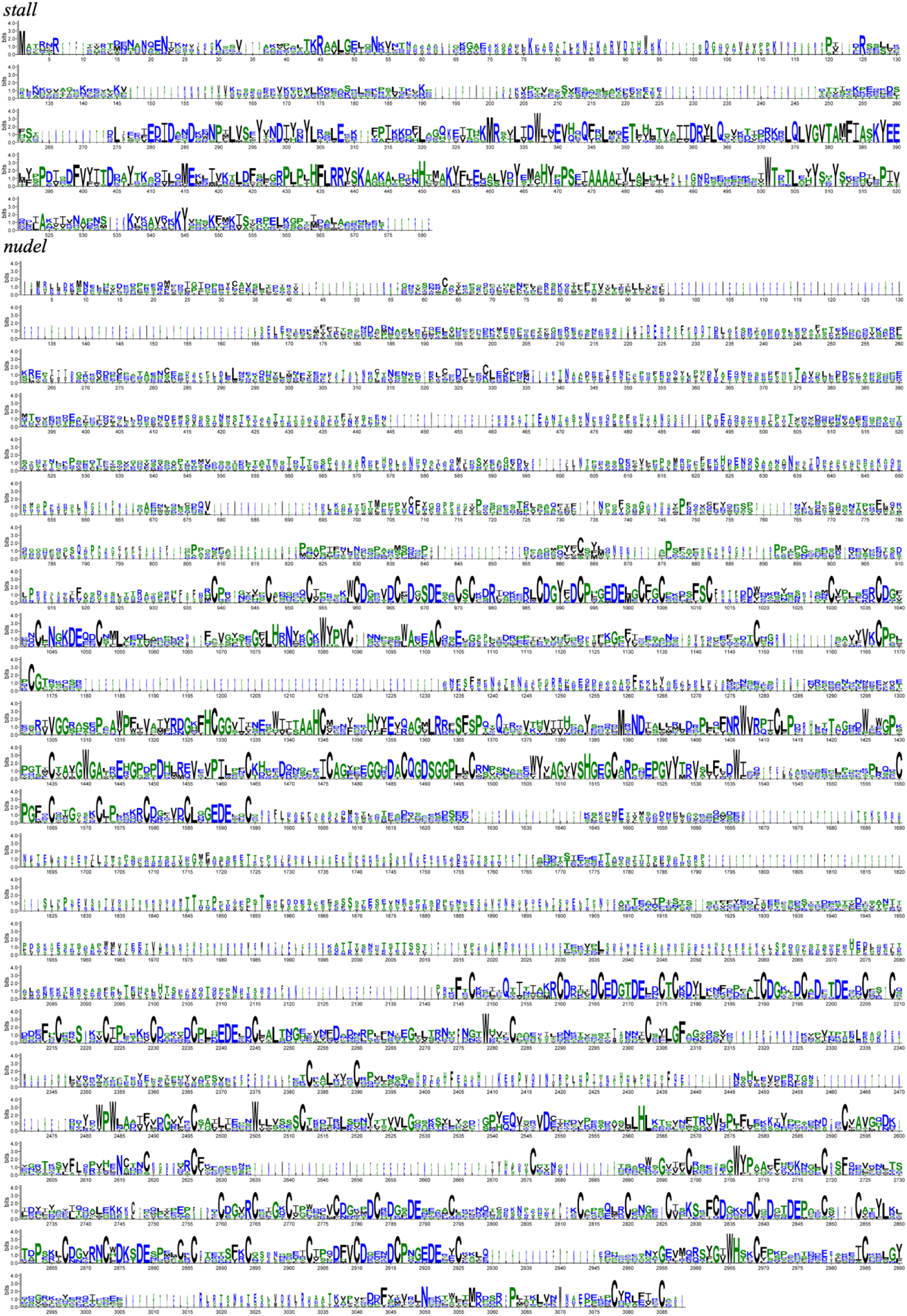
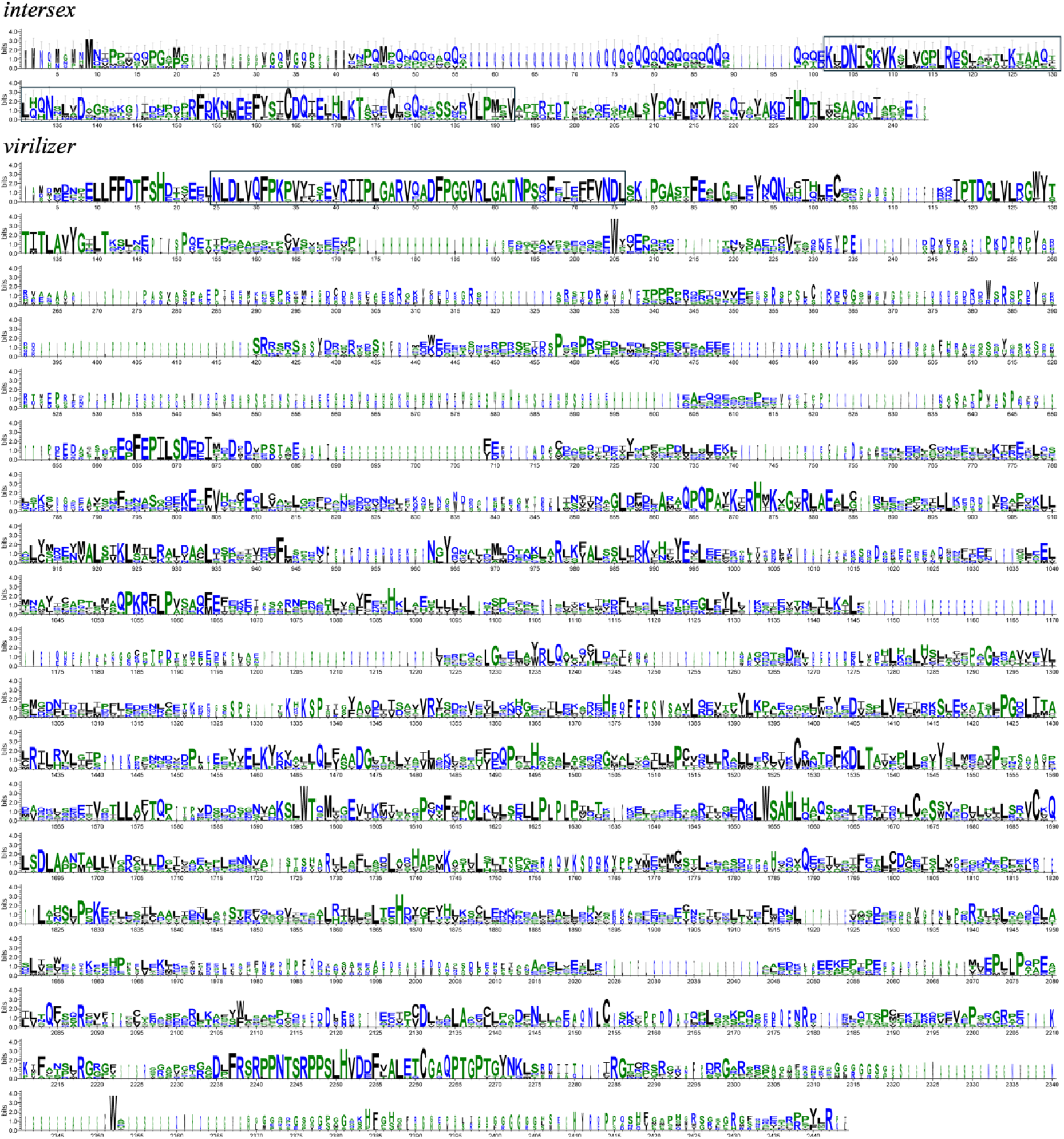
Multiple sequence alignment of candidate female fertility genes. Alignment includes sequences from *Solenopsis Invicta*, *Linepithema humile*, *Aedes aegypti*, *Anopheles stephensi*, *Drosophila melanogaster*, *Vespula vulgaris*, *Bemisia tabaci* and *Spodoptera frugiperda*. The region marked with a square is highly conserved at the animo acid level and was used for gRNA design (see Figure 11). Sequences for alignment were downloaded from the NCBI reference database. The online platform WebLogo (https://weblogo.berkeley.edu/logo.cgi) was used to visualize multiple protein sequence alignment. See next page for *stall* and *nudel* genes.

## References

1. Taber, S. W. Fire Ants. (Texas A&M University Press, 2000).

2. Wilson, E. O. Origin of the Variation in the Imported Fire Ant. Evolution 7, 262–263 (1953).

3. Fadamiro, H. Y., He, X. & Chen, L. Aggression in imported fire ants: an explanation for shifts in their spatial distributions in Southern United States? Ecol. Entomol. 34, 427–436 (2009).

4. Callcott, A.-M. A. & Collins, H. L. Invasion and Range Expansion of Imported Fire Ants (Hymenoptera: Formicidae) in North America from 1918-1995. Fla. Entomol. 240–240 (1996).

5. Chen, J. S. C., Shen, C.-H. & Lee, H.-J. Monogynous and Polygynous Red Imported Fire Ants, Solenopsis invicta Buren (Hymenoptera: Formicidae), in Taiwan. Environ. Entomol. 35, 167–172 (2006).

6. Wylie, R., Jennings, C., McNaught, M. K., Oakey, J. & Harris, E. J. Eradication of two incursions of the Red Imported Fire Ant in Queensland, Australia. doi:10.1111/emr.12197.

7. Moloney, S. & Vanderwoude, C. Red Imported Fire Ants: A threat to eastern Australia’s wildlife? Ecol. Manag. Restor. 3, 167–175 (2002).

8. Wylie, R., Yang, C.-C. S. & Tsuji, K. Invader at the gate: The status of red imported fire ant in Australia and Asia. Ecol. Res. 35, 6–16 (2020).

9. Red fire ants, a dreaded pest, have invaded Europe. https://www.science.org/content/article/red-fire-ants-dreaded-pest-have-invaded-europe.

10. Morrison, L. W. Long-term impacts of an arthropod-community invasion by the imported fire ant, solenopsis invicta. Ecology 83, 2337–2345 (2002).

11. Porter, S. D. & Savignano, D. A. Invasion of Polygyne Fire Ants Decimates Native Ants and Disrupts Arthropod Community. Ecology 71, 2095–2106 (1990).

12. Lard, C., et al. An Economic Impact of Imported Fire Ants in the United States of America. (2006).

13. Wylie, F. R. & Janssen-May, S. Red Imported Fire Ant in Australia: What if we lose the war? Ecol. Manag. Restor. 18, 32–44 (2017).

14. deShazo, R. D., Griffing, C., Kwan, T. H., Banks, W. A. & Dvorak, H. F. Dermal hypersensitivity reactions to imported fire ants. J. Allergy Clin. Immunol. 74, 841–847 (1984).

15. deShazo, R. D., Butcher, B. T. & Banks, W. A. Reactions to the Stings of the Imported Fire Ant. The New England Journal of Medicine https://www.nejm.org/doi/full/10.1056/NEJM199008163230707 (1990) doi:10.1056/NEJM199008163230707.

16. deShazo, R. D. & Banks, W. A. Medical consequences of multiple fire ant stings occurring indoors. J. Allergy Clin. Immunol. 93, 847–850 (1994).

17. Hays, S. B. & Arant, F. S. Insecticidal Baits for Control of the Imported Fire Ant, Solenopsis saevissima richteri1. *J. Econ. Entomol.* 53, 188–191 (1960).

18. Lofgren, C. S., Banks, W. A. & Glancey, B. M. Biology and Control of Imported Fire Ants. Annu. Rev. Entomol. 20, 1–30 (1975).

19. Burt, A. & Crisanti, A. Gene drive: Evolved and synthetic. *ACS Chem. Biol.* 13, 343–346 (2018).

20. Dhole, S., Lloyd, A. L. & Gould, F. Gene drive dynamics in natural populations: The importance of density dependence, space, and sex. Annu. Rev. Ecol. Evol. Syst. 51, 505–531 (2020).

21. Champer, J., Buchman, A. & Akbari, O. S. Cheating evolution: engineering gene drives to manipulate the fate of wild populations. Nat Rev Genet 17, 146–159 (2016).

22. Du, J. et al. Germline Cas9 promoters with improved performance for homing gene drive. Nat. Commun. 15, 4560 (2024).

23. Yadav, A. K., et al. CRISPR/Cas9-based split homing gene drive targeting doublesex for population suppression of the global fruit pest Drosophila suzukii. Proc. Natl. Acad. Sci. 120, e2301525120 (2023).

24. Kyrou, K. et al. A CRISPR-Cas9 gene drive targeting doublesex causes complete population suppression in caged Anopheles gambiae mosquitoes. Nat. Biotechnol. 36, 1062– 1066 (2018).

25. Meccariello, A. et al. Gene drive and genetic sex conversion in the global agricultural pest Ceratitis capitata. Nat. Commun. 15, 372 (2024).

26. Teem, J. L. et al. Genetic biocontrol for invasive species. Front. Bioeng. Biotechnol. 8, 452 (2020).

27. de la Filia, A. G., Bain, S. A. & Ross, L. Haplodiploidy and the reproductive ecology of Arthropods. Curr. Opin. Insect Sci. 9, 36–43 (2015).

28. Champer, J., Kim, I. K., Champer, S. E., Clark, A. G. & Messer, P. W. Suppression gene drive in continuous space can result in unstable persistence of both drive and wild-type alleles. Mol. Ecol. 30, 1086–1101 (2021).

29. Li, J. et al. Can CRISPR gene drive work in pest and beneficial haplodiploid species? Evol. Appl. 13, 2392–2403 (2020).

30. Liu, Y. & Champer, J. Modelling homing suppression gene drive in haplodiploid organisms. Proc. R. Soc. B 289, (2022).

31. Keller, L. The Assessment of Reproductive Success of Queens in Ants and Other Social Insects. Oikos 67, 177–180 (1993).

32. Keller, L. Social life: the paradox of multiple-queen colonies. Trends Ecol. Evol. 10, 355–360 (1995).

33. Obin, M. S., Morel, L. & Vander Meer, R. K. Unexpected, well-developed nestmate recognition in laboratory colonies of polygyne imported fire ants (Hymenoptera: Formicidae). J. Insect Behav. 6, 579–589 (1993).

34. Ross, K. G. Multilocus Evolution in Fire Ants: Effects of Selection, Gene Flow and Recombination. Genetics 145, 961–974 (1997).

35. Huang, Y.-C. & Wang, J. Did the fire ant supergene evolve selfishly or socially?: Prospects & Overviews. BioEssays 36, 200–208 (2014).

36. Macom, T. E. & Porter, S. D. Comparison of Polygyne and Monogyne Red Imported Fire Ant (Hymenoptera: Formicidae) Population Densities. Ann. Entomol. Soc. Am. 89, 535– 543 (1996).

37. Bull, J. J., Remien, C. H. & Krone, S. M. Gene-drive-mediated extinction is thwarted by population structure and evolution of sib mating. Evol. Med. Public Health 2019, 66–81 (2019).

38. North, A. R., Burt, A. & Godfray, H. C. J. Modelling the potential of genetic control of malaria mosquitoes at national scale. BMC Biol. 17, 26 (2019).

39. North, A. R., Burt, A. & Godfray, H. C. J. Modelling the suppression of a malaria vector using a CRISPR-Cas9 gene drive to reduce female fertility. BMC Biol. 18, 98 (2020).

40. Liu, Y., Teo, W., Yang, H. & Champer, J. Adversarial interspecies relationships facilitate population suppression by gene drive in spatially explicit models. Ecol. Lett. 26, 1174–1185 (2023).

41. Kim, J. et al. Incorporating ecology into gene drive modelling. Ecol. Lett. 26, S62–S80 (2023).

42. Chen, W., Guo, J., Liu, Y. & Champer, J. Population suppression by release of insects carrying a dominant sterile homing gene drive targeting doublesex in Drosophila. Nat. Commun. 15, 8053 (2024).

43. Faber, N. R. et al. Improving the suppressive power of homing gene drive by co-targeting a distant-site female fertility gene. Nat. Commun. 15, 9249 (2024).

44. Champer, S. E. et al. Computational and experimental performance of CRISPR homing gene drive strategies with multiplexed gRNAs. Sci. Adv. 6, eaaz0525 (2020).

45. Yang, E. et al. A homing suppression gene drive with multiplexed gRNAs maintains high drive conversion efficiency and avoids functional resistance alleles. G3 GenesGenomesGenetics (2022) doi:10.1093/G3JOURNAL/JKAC081.

46. Haller, B. C. & Messer, P. W. SLiM 4: Multispecies Eco-Evolutionary Modeling. Am. Nat. (2023) doi:10.1086/723601.

47. Vargo, E. L. & Fletcher, D. J. C. Effect of queen number on the production of sexuals in natural populations of the fire ant, Solenopsis invicta. Physiol. Entomol. 12, 109–116 (1987).

48. Fritz, G. N., Vander Meer, R. K. & Preston, C. A. Selective Male Mortality in the Red Imported Fire Ant, Solenopsis invicta. Genetics 173, 207–213 (2006).

49. Dhami, M. K. & Booth, K. Review of dispersal distances and landing site behaviour of Solenopsis invicta Buren, Red Imported Fire Ant (RIFA).

50. Goodisman, M. A. D., DeHeer, C. J. & Ross, K. G. Unusual Behavior of Polygyne Fire Ant Queens on Nuptial Flights. J. Insect Behav. (2000).

51. Bargum, K., Helanterä, H. & Sundström, L. Genetic population structure, queen supersedure and social polymorphism in a social Hymenoptera. J. Evol. Biol. 20, 1351–1360 (2007).

52. Keller, L. & Genoud, M. Extraordinary lifespans in ants: a test of evolutionary theories of ageing. Nature 389, 958–960 (1997).

53. Goodisman, M. A. D. & Ross, K. G. Queen recruitment in a multiple-queen population of the fire ant Solenopsis invicta. Behav. Ecol. 10, 428–435 (1999).

54. Ross, K. G., Vargo, E. L. & Keller, L. Social evolution in a new environment: the case of introduced fire ants. Proc. Natl. Acad. Sci. 93, 3021–3025 (1996).

55. Michel Chapuisat. Supergenes as drivers of ant evolution. (2023) doi:10.25849/MYRMECOL.NEWS_033:001.

56. Fire Ant Queen Longevity and Age: Estimation by Sperm Depletion | Annals of the Entomological Society of America | Oxford Academic. https://academic.oup.com/aesa/article/80/2/263/128254.

57. Markin, G. P., Dillier, J. H. & Collins, H. L. Growth and Development of Colonies of the Red Imported Fire Ant, Solenopsis invicta1. Ann. Entomol. Soc. Am. 66, 803–808 (1973).

58. Tschinkel, W. R. Colony growth and the ontogeny of worker polymorphism in the fire ant, Solenopsis invicta. Behav. Ecol. Sociobiol. 22, 103–115 (1988).

59. Battles between ants (Hymenoptera: Formicidae): a review. https://academic.oup.com/jinsectscience/article/24/3/25/7697941.

60. Greenberg, L., Vinson, S. B. & Ellison, S. Nine-Year Study of a Field Containing Both Monogyne and Polygyne Red Imported Fire Ants (Hymenoptera: Formicidae). Ann. Entomol. Soc. Am. 85, 686–695 (1992).

61. Meer, R. K. V., Cedeno, A. & Jaffe, K. Applied Myrmecology: A World Perspective. (CRC Press, 2019).

62. Kjeldgaard, M. K. et al. Distinct colony boundaries and larval discrimination in polygyne red imported fire ants (Solenopsis invicta). Mol. Ecol. 31, 1007–1020 (2022).

63. Wang, L., Xu, Y., Zeng, L. & Lu, Y. Impact of the red imported fire ant Solenopsis invicta Buren on biodiversity in South China: A review. J. Integr. Agric. 18, 788–796 (2019).

64. Fritz, G. N. & Vander Meer, R. K. Sympatry of Polygyne and Monogyne Colonies of the Fire Ant Solenopsis invicta (Hymenoptera: Formicidae). Ann. Entomol. Soc. Am. 96, 86–92 (2003).

65. Lin, C., Wen, T., Liu, Y., Huang, R. & Liu, H. K. Elucidating how the red imported fire ant (Solenopsis invicta) diffused spatiotemporally among different landscapes in north Taiwan, 2008–2015. Ecol. Evol. 11, 18604–18614 (2021).

66. Morianou, I. et al. Engineering Resilient Gene Drives Towards Sustainable Malaria Control: Predicting, Testing and Overcoming Target Site Resistance. 2024.10.21.618489 Preprint at 10.1101/2024.10.21.618489 (2024).

67. Adams, E. S. & Tschinkel, W. R. Mechanisms of population regulation in the fire ant Solenopsis invicta: an experimental study. J. Anim. Ecol. 70, 355–369 (2001).

68. Edgar, R. C. MUSCLE: multiple sequence alignment with high accuracy and high throughput. Nucleic Acids Res. 32, 1792–1797 (2004).

69. Jalview Version 2—a multiple sequence alignment editor and analysis workbench | Bioinformatics | Oxford Academic. https://academic.oup.com/bioinformatics/article/25/9/1189/203460.

70. Liu, Y. & Champer, J. Modelling homing suppression gene drive in haplodiploid organisms. Proc. R. Soc. B Biol. Sci. 289, 20220320 (2022).

71. Olejarz, J. W. & Nowak, M. A. Gene drives for the extinction of wild metapopulations. J. Theor. Biol. 577, 111654 (2024).

72. Champer, S. E., Kim, I. K., Clark, A. G., Messer, P. W. & Champer, J. Anopheles homing suppression drive candidates exhibit unexpected performance differences in simulations with spatial structure. eLife 11, (2022).

73. Zhang, X., Sun, W., Kim, I. K., Messer, P. W. & Champer, J. Population dynamics in spatial suppression gene drive models and the effect of resistance, density dependence, and life history. bioRxiv 2024.08.14.607913 (2024) doi:10.1101/2024.08.14.607913.

74. Xu, X. et al. Assessing target genes for homing suppression gene drive. 2024.12.06.627146 Preprint at 10.1101/2024.12.06.627146 (2024).

75. Verma, P., Reeves, R. G., Simon, S., Otto, M. & Gokhale, C. S. The Effect of Mating Complexity on Gene Drive Dynamics. Am. Nat. 201, E1–E22 (2023).

76. Shoemaker, D. D., Costa, J. T. & Ross, K. G. Estimates of heterozygosity in two social insects using a large number of electrophoretic markers. Heredity 69, 573–582 (1992).

77. Ross, K. G. & Fletcher, D. J. C. Comparative study of genetic and social structure in two forms of the fire ant Solenopsis invicta (Hymenoptera: Formicidae). Behav. Ecol. Sociobiol. 17, 349–356 (1985).

78. Porter, S. D. Stability of Polygyne and Monogyne Fire Ant Populations (Hymenoptera: Formicidae: Solenopsis invicta) in the United States. J. Econ. Entomol. 86, 1344–1347 (1993).

79. Yang, C.-C. S. et al. Successful establishment of the invasive fire ant Solenopsis invicta in Taiwan: insights into interactions of alternate social forms. Divers. Distrib. 15, 709–719 (2009).

80. On the relationship between queen number and fecundity in polygyne colonies of the fire ant Solenopsis invicta - VARGO - 1989 - Physiological Entomology - Wiley Online Library. https://resjournals.onlinelibrary.wiley.com/doi/abs/10.1111/j.1365-3032.1989.tb00955.x?casa_token=3HGFCG5gcJIAAAAA:xIZdO8Gs8D9CZ6OekRfxIb9R79IMraUtiJmlF3mAYMDBhW8KU736iSwNxHz82qC0Yakxskyz7ZdgVKv6KQ.

81. Gardner, A. & West, S. A. Greenbeards. Evolution 64, 25–38 (1,January).

82. Blackburn, T. M. et al. A proposed unified framework for biological invasions. Trends Ecol. Evol. 26, 333–339 (2011).

83. Chapple, D. G. et al. Biological invasions as a selective filter driving behavioral divergence. Nat. Commun. 13, 5996 (2022).

84. Les, G. Encyclopedia of Insects. (2009).

85. Native Ants. Texas Imported Fire Ant Research and Management Project https://fireant.tamu.edu/learn/native-ants/.

86. Adams, E. S. Experimental analysis of territory size in a population of the fire ant Solenopsis invicta. Behav. Ecol. 14, 48–53 (2003).

87. Tschinkel, W. R. Brood raiding and the population dynamics of founding and incipient colonies of the fire ant, Solenopsis invicta. Ecol. Entomol. 17, 179–188 (1992).

88. Chiu, Y.-K., Hsu, J.-C., Chang, T., Huang, Y.-C. & Wang, J. Mutagenesis mediated by CRISPR/Cas9 in the red imported fire ant, Solenopsis invicta. Insectes Sociaux 67, 317–326 (2020).

89. Hart, T. et al. Sparse and stereotyped encoding implicates a core glomerulus for ant alarm behavior. Cell 186, 3079–3094.e17 (2023).

